# Joint Modeling of Gene-Environment Correlations and Interactions using Polygenic Risk Scores in Case-Control Studies

**DOI:** 10.1101/2023.02.14.528572

**Authors:** Ziqiao Wang, Wen Shi, Raymond J. Carroll, Nilanjan Chatterjee

## Abstract

Polygenic risk scores (PRS) are rapidly emerging as aggregated measures of disease-risk associated with many genetic variants. Understanding the interplay of PRS with environmental factors is critical for interpreting and applying PRS in a wide variety of settings. We develop an efficient method for simultaneously modeling gene-environment correlations and interactions using PRS in case-control studies. We use a logistic-normal regression modeling framework to specify the disease risk and PRS distribution in the underlying population and propose joint inference across the two models using the retrospective likelihood of the case-control data. Extensive simulation studies demonstrate the flexibility of the method in trading-off bias and efficiency for the estimation of various model parameters compared to the standard logistic regression or a case-only analysis for gene-environment interactions, or a control-only analysis for gene-environment correlations. Finally, using simulated case-control datasets within the UK Biobank study, we demonstrate the power of the proposed method for its ability to recover results from the full prospective cohort for the detection of an interaction between long-term oral contraceptive use and PRS on the risk of breast cancer. This method is computationally efficient and implemented in a user-friendly R package.

## 1 Introduction

Large-scale genome-wide association studies (GWAS) are now leading the development of polygenic risk scores (PRS), also known as polygenic scores (PGS), as a measure of the total genetic burden individuals carry across many genetic variants associated with different traits and diseases (Khera et al., 2018; Torkamani et al., 2018; Lewis and Vassos, 2020; Kullo et al., 2022). As the sample size for GWAS continues to increase, PRS are becoming increasingly predictive of underlying complex traits. There are growing interests about their applications in a number of different contexts, including but not limited to risk stratification (Chatterjee et al., 2016), improving the accuracy of diagnostic biomarkers (Kachuri et al., 2022), and as research tools for understanding genetic links between traits (Dennis et al., 2021). Applications of PRS in the future, however, will require better characterization of its interplay with environmental variables, including societal, lifestyle, and behavioral factors.

Gene-environment interplay can manifest in two forms: correlations and interactions. PRS for a given disease may show a correlation with an environmental factor, which has a heritable component itself, due to systematic pleiotropic associations of underlying variants across multiple traits. PRS can also exhibit correlation with environmental factors, such as social determinants of health, due to the presence of underlying population structure across which the distribution of both genes and environments vary (Haworth et al., 2019; Zaidi and Mathieson, 2020). Gene-environment interaction, on the other hand, typically refers to the non-additive or non-multiplicative effects of the two factors on a given trait. Modeling gene-environment interactions using PRS can lead to improved power when individual SNPs have weak effects but they collectively follow a common pattern of interactions (Domingue et al., 2020; Mas et al., 2020; Blechter et al., 2021; Jacobs et al., 2021). PRS can also be used to test for interaction of an exposure with a group of genetic variants restricted to some relevant gene sets, like biological pathways, that may be mediating the effect of the exposures. Further, as GWAS-derived PRS are now getting primed to be widely used for risk prediction, it is increasingly imperative to characterize their interactions with other risk factors of corresponding diseases for the development of comprehensive risk-prediction model including both genetic and non-genetic factors (Chatterjee et al., 2016).

Case-control studies, because of their time and economic efficiency, remain widely used for conducting epidemiologic studies of rare diseases. Standard logistic regression analysis is commonly used for statistical inference on disease odd-ratio parameters, including those of the interaction terms, from the analysis of case-control studies. Further, controls from case-control studies are also commonly used to describe characteristics of risk factors and their interrelationships in the underlying population assuming that the disease is rare. There exists, however, opportunities for more efficient inference in case-control studies exploiting a retrospective likelihood of the data. In particular, for the analysis of gene-environment interactions using single-SNP genotype data, a variety of methods (Piegorsch et al., 1994; Umbach and Weinberg, 1997; Chatterjee and Carroll, 2005; Mukherjee and Chatterjee, 2008; Murcray et al., 2009; Hsu et al., 2012; Gauderman et al., 2017) have been developed to explore an assumption of gene-environment independence in the underlying population to enhance the efficiency of inference on interaction odd-ratio parameters. Conversely, it has been shown that gene-environment correlations can be estimated efficiently by pooling cases and controls from a case-control study in the absence of gene-environment interactions (Li et al., 2010). We have earlier introduced a case-only method for studying gene-environment interactions using PRS (Meisner et al., 2019), but the method is limited because of its inability to leverage data on controls. It cannot provide estimates of a full spectrum of model parameters, including the main effects of the environmental variables in an underlying logistic model or gene-environment correlation parameters in the population.

In this study, we propose a joint modeling approach to gene-environment interactions and correlations using PRS based on a logistic-normal framework. The logistic model for disease risk can be arbitrarily flexible to include the main effects of PRS, environmental variables, factors related to population stratification, and other confounders, and any relevant interaction terms. Similarly, the normal model for the PRS distribution can be arbitrarily flexible to allow the mean of PRS in the underlying population to vary by a combination of different user-specified factors, including categorical and continuous markers of population stratification, and environmental factors. Further, heteroscedasticity in PRS distribution is allowed according to a categorical variable, such as self-reported ancestry or country of origin. We then use a profile-retrospective-likelihood technique (Chatterjee and Carroll, 2005) for developing a robust and speedy computational algorithm for the joint estimation of all of the parameters across the two models.

We use simulated data to compare the performance of the proposed method to several alternative methods, including standard logistic regression, a case-only method (Meisner et al., 2019), and a nonparametric regression approach (Stalder et al., 2017) for the estimation of gene-environment interactions; and control-only analysis, weighted and adaptively weighted estimators (Li et al., 2010) for the estimation of gene-environment correlations. We further simulate case-control studies for incident breast and colorectal cancers from the UK Biobank study and apply the proposed method to evaluate its ability to recover results from the analysis of the underlying full cohort. We discover an interaction effect between long-term oral contraceptive (OC) use and PRS on the risk of postmenopausal and premenopausal breast cancer, respectively; and an interaction effect between age and PRS on the risk of colorectal cancer. Both simulation studies and data applications demonstrate the utility of the method for efficient and flexible analysis of PRS by *E* correlations and interactions in case-control studies. Our method is implemented in R, with a standard syntax as other popularly used regression modeling functions.

## 2 Material and Methods

### 2.1 Model Specification and Assumptions

Let *D* denote a binary outcome variable with values 0 and 1, indicating the status of a disease. Let the marginal prevalence of the disease in the underlying population be denoted by *π*_1_ = pr(*D* = 1). Let *Z* denote the PRS value and ***E*** denote a vector of environmental variables of interest. Additionally, we let ***S*** be a potential vector-valued co-factors that may influence the distribution of PRS in the underlying population. ***S*** may, for example, include some components of ***E*** itself which may be suspected to be correlated with PRS, such as factors related to social determinants of health or environmental factors with strong heritable components, and population stratification factors, such as self-reported race/ethnicity and genetic principal components (PCs).

We assume pr(*Z*|***E, S***) = pr(*Z*|***S***), i.e., Z is independent of ***E*** conditional on ***S***. We further define

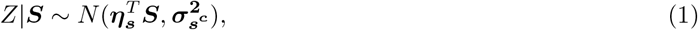

where in the mean component, ***η***_***s***_ denotes a set of unknown parameters characterizing the linear correlation between *Z* and components of ***S***, and the variance of the normal distribution is allowed to vary by the levels of a categorical stratification variable *S*^*c*^, assumed to be a coarsened version of ***S***. As PRS is typically defined by a combination of many genetic variants, the normality approximation is typically accurate due to the central limit theorem. Further, in our software, users have the complete flexibility to specify the mean model and embed any assumption of gene-environment independence or dependence by the exclusion or inclusion of the corresponding ***E/S*** variables. Moreover, we allow for heteroscedasticity in the distribution of PRS, but here instead of specifying a regression model, we allow the variance parameters to freely vary according to a user-specified categorical variable. In general, ***S*** can include continuous and categorical variables, such as the top 10 PCs, self-reported race/ethnic groups, and geographic regions; and *S*^*c*^ can be constructed by one or a combination of multiple categorical variables such as race/ethnic groups and geographic regions.

Next, we define the disease model of interest in the form

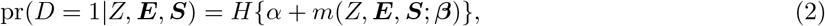

where *H*(*x*) = *{*1 + exp(−*x*)*}*^−1^ is the logistic distribution function and *m*(·) is a known but arbitrary function. In the typical logistic regression model, the function *m*(·) is assumed to be linear in parameters, with the resulting scale corresponding to the linear logistic but a non-linear form for *m*(·) can potentially be considered if interactions were to be modeled in alternative scales (Chatterjee and Carroll, 2005; Han et al., 2012). In this article, we let *H{α* + *m*(*Z*, ***E, S***; ***β***)*}* follow a standard logistic form with *α* being the log odds of the disease in a baseline category and ***β*** being the log odds ratio parameters associated with different risk factors and their interactions.

### 2.2 The Case-Control Sampling Design and Retrospective Profile Likelihood

We assume that under a case-control design, *N*_1_ cases and *N*_0_ controls are sampled from an underlying population for which the above models hold. The retrospective likelihood for case-control data takes the form 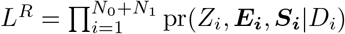, where, using Bayes theorem, each individual term could be derived in the form

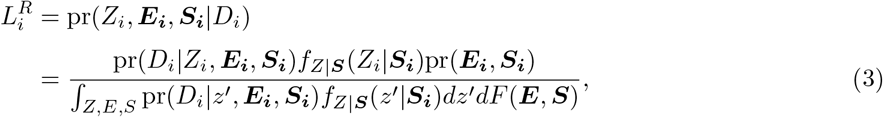

where the joint distribution function for (***E, S***) remains completely unspecified.

The maximum likelihood estimates for the parameters of interest, namely ***β, η***_***s***_, and 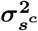 can be obtained using the profile-likelihood techniques as described in Chatterjee and Carroll (2005). We derive the form of the profile likelihood in the general case where we consider the interaction effects between *Z* and ***X*** = (***E, S***), with

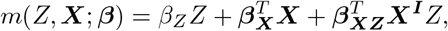

where 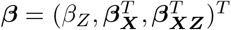, with ***X***^***I***^ ⊆ ***X*** involves the environmental and stratification variables for which interactions are to be incorporated in relationship to the PRS variable.

We show in Appendix A that a profile-likelihood of the data, which removes high-dimensional nuisance parameters associated with the distribution of pr(***E, S***), can be approximated under a rare disease assumption as

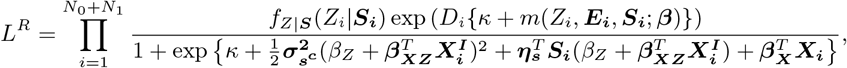

where we define a nuisance intercept *κ* = *α* + *log*(*N*_1_*/N*_0_) − *log*(*π*_1_*/π*_0_). We observe that the intercept *α* in the disease model (2) can be estimated with the above likelihood only if the disease prevalence *π*_1_ is known *a priori*. In a situation of frequency-matched case-control design, the proposed method can be easily modified by defining intercept terms 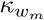 for each of the *M* strata in the matching variables *W* = *w*_*m*_(*m* = 1, · · ·, *M*). Details of the derivation and the log-likelihood score function are provided in Appendix A. The maximum likelihood estimation for the unknown parameters 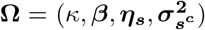 can be efficiently calculated with the input of score function using quasi-Newton algorithm within a few iterations.

### 2.3 Simulation Studies

We conducted extensive simulation studies to evaluate the bias, efficiency, and type I errors associated with various parameters of interest underlying the models. We first simulated a cohort of 10^6^ individuals with a population disease prevalence of *π* = Pr(*D* = 1) = 0.01 using a disease model assuming a logistic regression model of the form:

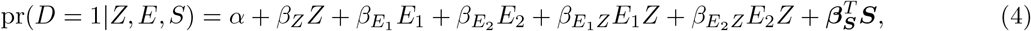

where *Z* denotes the PRS value; *E*_1_, a binary variable, and *E*_2_, a continuous variable, denote two environmental variables that are independent of *Z*; ***S*** = (*S*_1_, *S*_2_) is a vector of stratification variables that include a categorical variable *S*_1_ of three levels and a continuous variable *S*_2_ that are both correlated with *Z*. Using the above model, we consider a range of scenarios of gene-environment correlations and interactions (see scenarios in Table 1). We applied our method to a randomly selected *N*_0_ = *N*_1_ = 500 case-control study from the underlying cohort, each with 1 000 simulations. Detailed descriptions of the parameter values are in Appendix B Section 2.1.

**Table 1:** Model and parameter setup for the 6 scenarios in the simulation study. Scenario 1 evaluates type I error of both interaction terms; scenarios 1,2,3, and 6 assume that all environmental variables are independent of each other; scenarios 4 and 5 assume that *S*_2_ and *E*_2_ follow a bivariate normal distribution with a correlation of 0.4. *S*_1_ is a categorical variable of 3 levels: *S*_1(1)_, *S*_2(1)_ are two dummy variables of *S*_1_. The true parameter values are 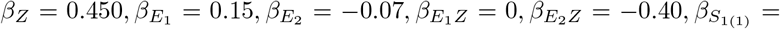 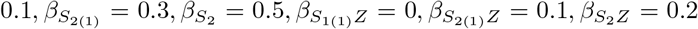, unless indicated in the notes. In the PRS model, 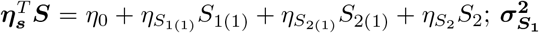 is a three-dimensional vector that corresponds to the 3 strata in *S*_1_. ***η***_***s***_ = *c*(1, 0.2, 0.3, 0.2) and 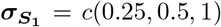 for each stratum in *S*_1_. The intercept term of the disease model is determined so that the population disease prevalence is fixed at around 0.01.

| Scenario | Disease Model: $\text{pr}(D = 1\vert Z, \mathbf{E}, \mathbf{S})$ | PRS Model: $Z\vert \mathbf{S}$ | Notes |
| --- | --- | --- | --- |
| 1 | $Z + E_1 + E_2 + E_1Z + E_2Z$ | $N(0, 1)$ | $\beta_{E_1Z} = 0, \beta_{E_2Z} = 0$ |
| 2 | $Z + E_1 + E_2 + S_{1(1)} + S_{2(1)} + S_2 + E_1Z + E_2Z$ | $N(\boldsymbol{\eta}_s^T \mathbf{S}, \boldsymbol{\sigma}_{\mathbf{S}_1}^2)$ | $\beta_{E_1Z} = 0$ |
| 3 | $Z + E_1 + E_2 + S_{1(1)} + S_{2(1)} + S_2 + E_1Z + E_2Z + S_{1(1)}Z + S_{2(1)}Z + S_2Z$ | $N(\boldsymbol{\eta}_s^T \mathbf{S}, \boldsymbol{\sigma}_{\mathbf{S}_1}^2)$ | $\beta_{E_1Z} = 0, \beta_{S_{1(1)}Z} = 0$ |
| 4 | $Z + E_1 + E_2 + E_1Z + E_2Z$ | $N(\boldsymbol{\eta}_s^T \mathbf{S}, \boldsymbol{\sigma}_{\mathbf{S}_1}^2)$ | $\beta_{E_1Z} = 0, \rho_{E_2, S_2} = 0.4$ |
| 5 | $Z + E_1 + E_2 + S_{1(1)} + S_{2(1)} + S_2 + E_1Z + E_2Z$ | $N(\boldsymbol{\eta}_s^T \mathbf{S}, \boldsymbol{\sigma}_{\mathbf{S}_1}^2)$ | $\beta_{E_1Z} = 0, \rho_{E_2, S_2} = 0.4$ |
| 6 | $Z + S_{1(1)} + S_{2(1)} + S_2 + S_{1(1)}Z + S_{2(1)}Z + S_2Z$ | $N(\boldsymbol{\eta}_s^T \mathbf{S}, \boldsymbol{\sigma}_{\mathbf{S}_1}^2)$ | $\beta_{S_{1(1)}Z} = 0$ |

To have a comprehensive understanding of the model performance, we compared our method to three existing methods for investigating PRS × *E* interaction: (1) the ordinary logistic regression, (2) a recently developed case-only method (Meisner et al., 2019), and (3) a nonparametric regression method (Stalder et al., 2017) that allows the distribution of PRS to remain unspecified. The case-only method requires simply fitting a linear regression model of the PRS on E and then scaling the estimates of regression coefficients suitably to estimate corresponding interaction parameters in an underlying logistic regression model. The method can further estimate the PRS main effect if supplemented with external data. There are two ways: one can use estimates of the mean and the standard deviation of the PRS for the underlying population from an external study, or one can import an external estimate of the mean but use the residual standard deviation from the internal case-only linear regression analysis itself to estimation the PRS population standard deviation. The method can also adjust for potential stratification factors, but it implicitly assumes that the underlying logistic regression model for the disease-risk incorporates interaction terms between PRS and the stratification factors. In other words, it does not have the flexibility of the traditional logistic regression analysis of specifying any disease model of choice. To implement the case-only method, we fitted linear regression models of the form *Z* ∼ *E*_1_ + *E*_2_ in scenarios 1 and 4; *Z* ∼ *E*_1_ + *E*_2_ + *S*_1(1)_ + *S*_2(1)_ + *S*_2_ in scenarios 2, 3, and 5; and *Z* ∼ *S*_1(1)_ + *S*_2(1)_ + *S*_2_ in scenario 6. The nonparametric method evaluates a semiparametric likelihood for case-control data similar to ours, but calculates underlying expectations with respect to PRS distribution using empirical mean as opposed to using normal moment generating functions. The method being nonparametric in PRS distribution, does not have the flexibility to incorporate the conditional distribution of [PRS|S] for complex sets of stratification factors as we allow in our modeling framework. We used an available software where in the disease model we incorporated ***E*** and ***S*** in the same way as our proposed method for scenarios 1, 3, 4, and 6, but the underlying assumption behind the nonparametric method is that PRS is independent of any environmental and stratification variable. Given that the software currently can only incorporate both main effect and interaction terms of environmental variables in the disease model, it over-adjusted for the interaction terms *Z* × ***S*** in scenarios 2 and 5. Both the case-only and nonparametric methods additionally rely on the rare disease assumption.

To assess the performance for the estimation of correlation parameters between PRS and environmental variables, we compared our method to an approach that uses linear regression based on the underlying PRS model *Z* ∼ *S*_1(1)_ + *S*_2(1)_ + *S*_2_ in control-only samples, a weighted estimator that integrates both controls and cases, and an adaptively weighted method (Li et al., 2010), details of the methods can be found in Appendix B Section 2.3.

### 2.4 Simulated Case-Control Study within UK Biobank

We applied the retrospective likelihood method to data from the UK Biobank, an ongoing large-scale prospective study of approximately 500 000 UK participants first recruited during 2006-2010 (Sudlow et al., 2015). We simulated two case-control studies within the UK Biobank, one for incident breast cancer and the other for incident colorectal cancer, and evaluated the performance of the proposed and alternative methods for the characterization of gene-environment interactions for these two malignancies. As data from the whole underlying cohort of the UK Biobank is available, we used results from an underlying logistic regression model for the full cohort to provide a benchmark for comparison of the methods for the analysis of the case-control datasets.

PRS for breast and colorectal cancers were derived using 313 and 95 variants respectively, based on weight available from the PGS catalog (Lambert et al., 2021; PGS Catalog Team, 2022) provided by recent external GWAS (Mavaddat et al., 2019; Huyghe et al., 2019). The incident cancer status for individuals was determined using cancer registry records available until June 2021. We removed individuals who were diagnosed with respective cancers before entry into the study. A total of 2 024 incident breast cancer cases and 89 241 controls for postmenopausal women; 704 cases and 37 785 controls for the premenopausal cohort in unrelated white female participants were analyzed. A total of 1 743 incident colorectal cancer cases and 282 429 controls in unrelated white individuals were analyzed. For each cancer, we selected all cases and a random sample of controls with equal size as the cases. The environmental variables (***E***) of interest for breast cancer were age, parity, age at first birth, alcohol intake, height, body mass index (BMI), age at menarche, years of OC use, age at menopause (for postmenopausal women), and hormone replacement therapy (HRT) use (for postmenopausal women). And those for colorectal cancer were age, gender, alcohol intake, height, waist circumference, smoking status, red meat intake, vegetable intake, processed meat intake per week, and activity levels. For both cancers, we additionally included the main effects of geographic birth locations and their interactions with PRS and adjusted for the top 10 genetic PCs and assessment centres as additional confounders in the disease model.

For the PRS models across both cancers, we make the assumption of independence between PRS and the environmental exposures conditional on ***S*** = (Top 10 PCs, North coordinates of birth location, East coordinates of birth location) and stratification factor *S*^*c*^ = (Assessment Centres). Thus the PRS model is specified as follows:

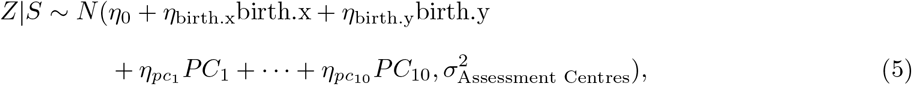

where birth.x and birth.y are the East and North coordinates of birth location respectively. Details of the analyses are available in Appendix C Sections 3.1 and 3.2.

## 3 Results

### 3.1 Simulation Studies

#### 3.1.1 Bias, Efficiency, and Type I Error in the Disease Model

Figure 1 shows the bias and efficiency of the PRS main effect and its interaction parameters in scenarios 1-6. Additional results on MSE of the estimates and on main effect estimates of environmental variables in the model can be found in Appendix B, Figure 1-5. We make a number of key observations from the simulation studies.

**Figure 1:**
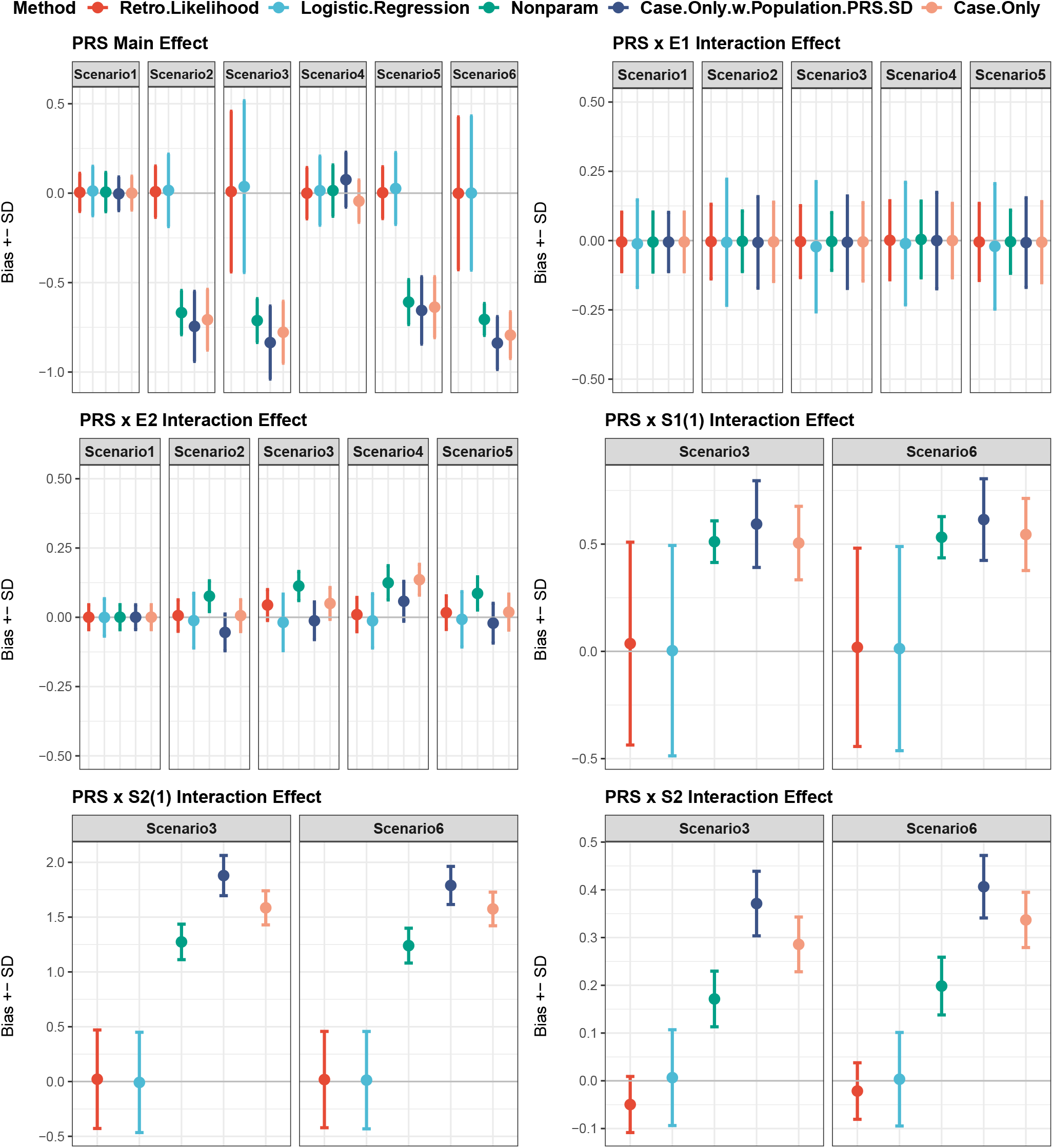
Performance of alternative methods for estimating parameters of the disease-risk model in simulation studies. The bias and estimated standard deviation (SD) of the PRS main effect log odds ratio *β*_*Z*_ in simulation scenarios 1-6; 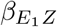 and 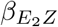 in scenarios 1-5; and 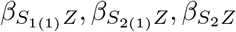 in scenario 3 and 6. In the figure, each dot represents bias and the error bar is SD.

First, between standard logistic regression and the retrospective-likelihood method, the latter method gained remarkable efficiency advantage for the estimation of PRS-***E*** interaction terms by exploiting the underlying independence assumption. For the estimation of PRS-***S*** interaction term, on the other hand, where the underlying PRS model accounted for PRS-***S*** correlation terms, the two methods performed similarly in terms of both bias and standard error. This shows the flexibility of the proposed method to trade-off bias and efficiency in estimation of G-E interaction parameters through flexible specification of the underlying G-E correlation model.

Second, the retrospective-likelihood method showed good bias-variance trade-off capacity compared to the case-only and nonparametric methods. In scenario 1, where there was no G-E or G-S correlations, the case-only and non-parametric methods performed similarly to the proposed retrospective likelihood approach in terms of maintaining unbiasedness and yet smaller standard errors compared to logistic regression. However, under scenarios 2-6, where there were G-S correlations, we observed that these methods tended to produce biased estimates for a number of terms, including PRS main effect (scenario 2,3, 5, and 6), interaction of PRS with *E*_2_ (scenario 2-6), and interaction of PRS with both of the stratification factors *S*_1_ and *S*_2_, but not for interaction of PRS with *E*_1_. The bias of these alternative methods stem from their inability to properly adjust for the correlation of PRS with the stratification factors ***S***. Overall, among all of the different methods, the proposed retrospective-likelihood approach generally maintained the smallest MSE for the estimation of different parameters due to its ability to trade-off bias and efficiency in a flexible manner (see Appendix B, Figures 1-3).

Finally, in terms of type I errors of the interaction terms, both the retrospective likelihood method and standard logistic regression maintained the nominal level in varying situations with different conditional independence assumptions between PRS and ***E***. The nonparametric and the case-only methods had highly inflated type I error for the stratification variable *S*_1_ as shown in scenarios 3 and 6 (Appendix B Table 1).

#### 3.1.2 Parameter Estimation of the PRS model

We investigated how our joint modeling approach estimated 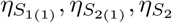 and 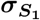 in the PRS model. Figure 2 summarizes the bias and estimated standard deviation for the regression coefficients ***η***_***s***_. For the binary variables *S*_1(1)_ and *S*_2(1)_, as the underlying disease model involved no interaction effect between PRS and *S*_1(1)_, our method had greater efficiency gain than other methods by utilizing both the cases and controls (shown in the top panels of Fig 2); when there was true interaction effect PRS x *S*_2(1)_, our method approximated the efficiency of the control-only method by only using the control samples in MLE (middle panels). Both weighted and adaptively weighted estimators were slightly more efficient than the proposed method estimating 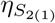 but tended to cause bias when the interaction effect was 0 because of weighing in the estimates using cases alone. The most efficiency gain was observed for the continuous variable *S*_2_ in the proposed method, regardless of whether the interaction term was 0 or not in the disease model (bottom panels). The retrospective likelihood method also produced unbiased estimates of PRS variance parameters, 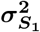 (Appendix B Table 2), further demonstrating the unique feature of explicating the heteroscedasticity of the population PRS values inherent from multiple stratification factors.

**Figure 2:**
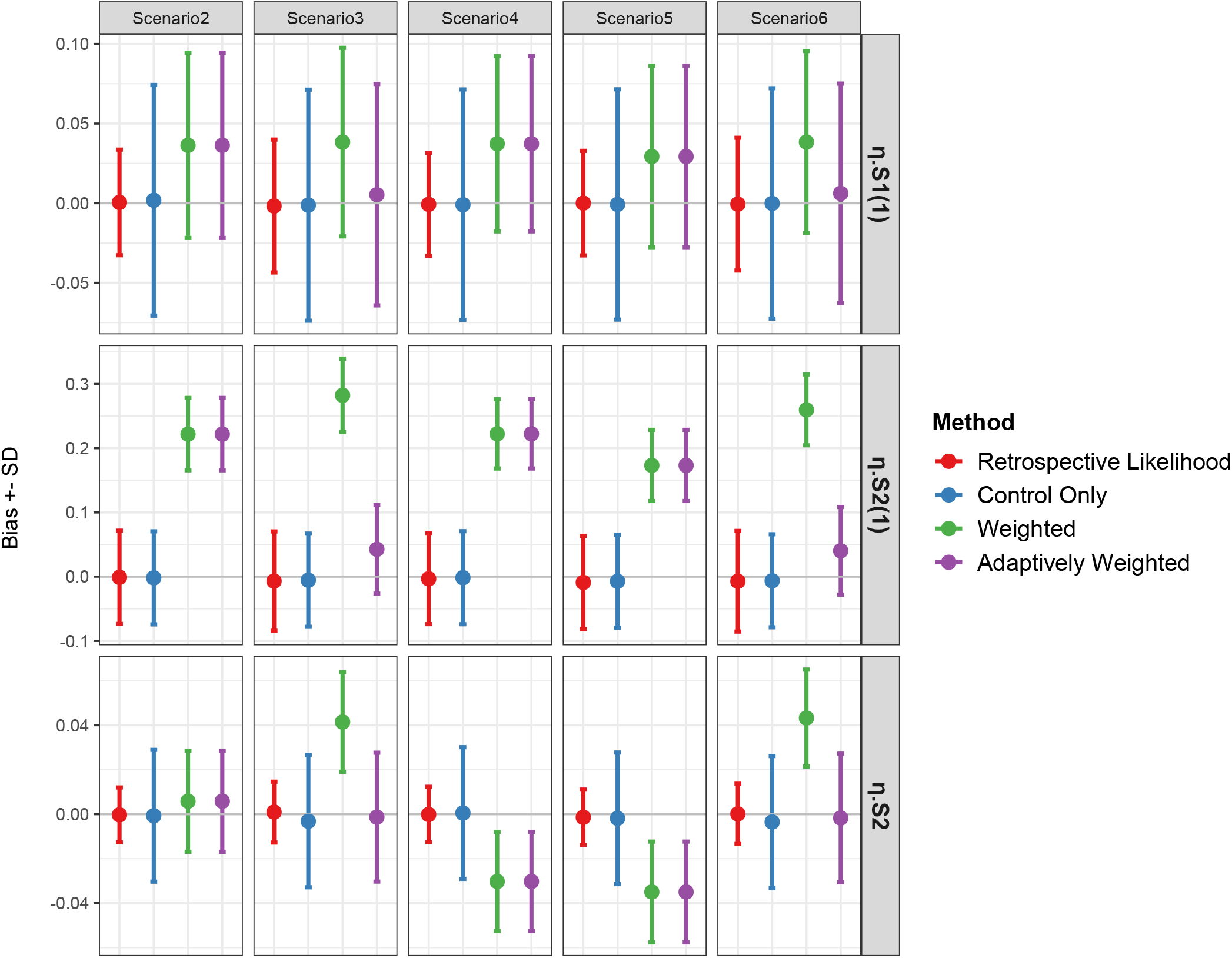
Performance of alternative methods for the estimation of PRS-S correlation parameters in simulation studies. The estimated regression coefficients 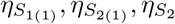, with true values (0.2, 0.3, 0.2) in the PRS model 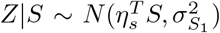 in simulation scenarios 2-6. The true interaction parameter values in the disease model are 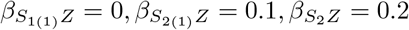.

### 3.2 Analysis of UK Biobank Incident Breast Cancer

In Table 2, we show the estimates of log odds ratio (log OR) for premenopausal and postmenopausal breast cancer with underlying PRS and two environmental variables, years of OC use and East coordinates of birth location. We selected these environmental variables to demonstrate the operational characteristics of the methods as they showed significant interaction with PRS in the underlying full cohort analysis. Results for the other parameters in the model can be found in Appendix C Tables 7 and 8. The logistic regression analysis of the full cohort data suggested a statistically significant interaction between PRS and long-term OC use for both postmenopausal and premenopausal women. We noticed similarly significant results from the retrospective likelihood analysis of the case-control data and the case-only analysis. However, this interaction effect was not significant in the ordinary logistic regression of the same set of case-control data. A significant efficiency gain of the retrospective-likelihood method over logistic regression was also observed for the estimation of PRS main effect. On the other hand, the case-only method, when supplemented with an estimate of the underlying population mean of the PRS, (Meisner et al., 2019), produced a more attenuated estimate of the PRS main effect compared to the full cohort analysis.

**Table 2:**
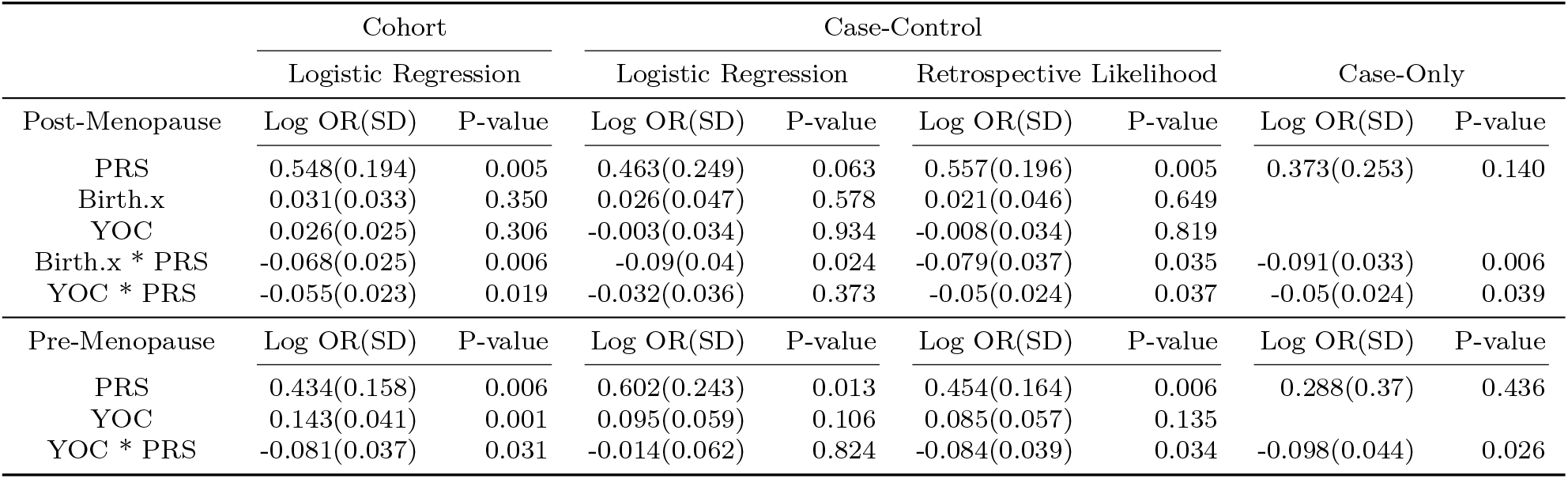
Parameter estimates of the log odds ratio (log OR), standard deviation, and p-values for the risk model in the UK Biobank breast cancer data for postmenopausal and premenopausal women respectively. PRS-associated risks are presented in terms of per-SD unit. YOC, years of oral contraceptive use; PRS, breast cancer-related polygenic risk score constructed from 313 SNPs; Birth.x, the East co-ordinates of individual’s birth location.

We additionally performed two sets of sensitivity analyses for the years of OC use by categorizing them into time intervals. The first test was to categorize the years of OC use into three categories: never, 0-10 years, and *>* 10 years. The second test contained more refined categories: never, 0-5 years, 5-10 years, 10-15 years, and *>* 15 years. In these analyses, we discovered a dose-response relationship in the interaction between the years of OC use and PRS for postmenopausal women in both the logistic regression analysis of the full prospective cohort and the retrospective-likelihood analysis of the case-control data but not in the logistic regression analysis of the case-control data (Figure 3 and Appendix C Table 3). We also observed a weak dose-response relationship when categorizing the years of OC use in the premenopausal cohort but the result was not statistically significant possibly due to lack of power (Appendix C Table 4).

**Figure 3:**
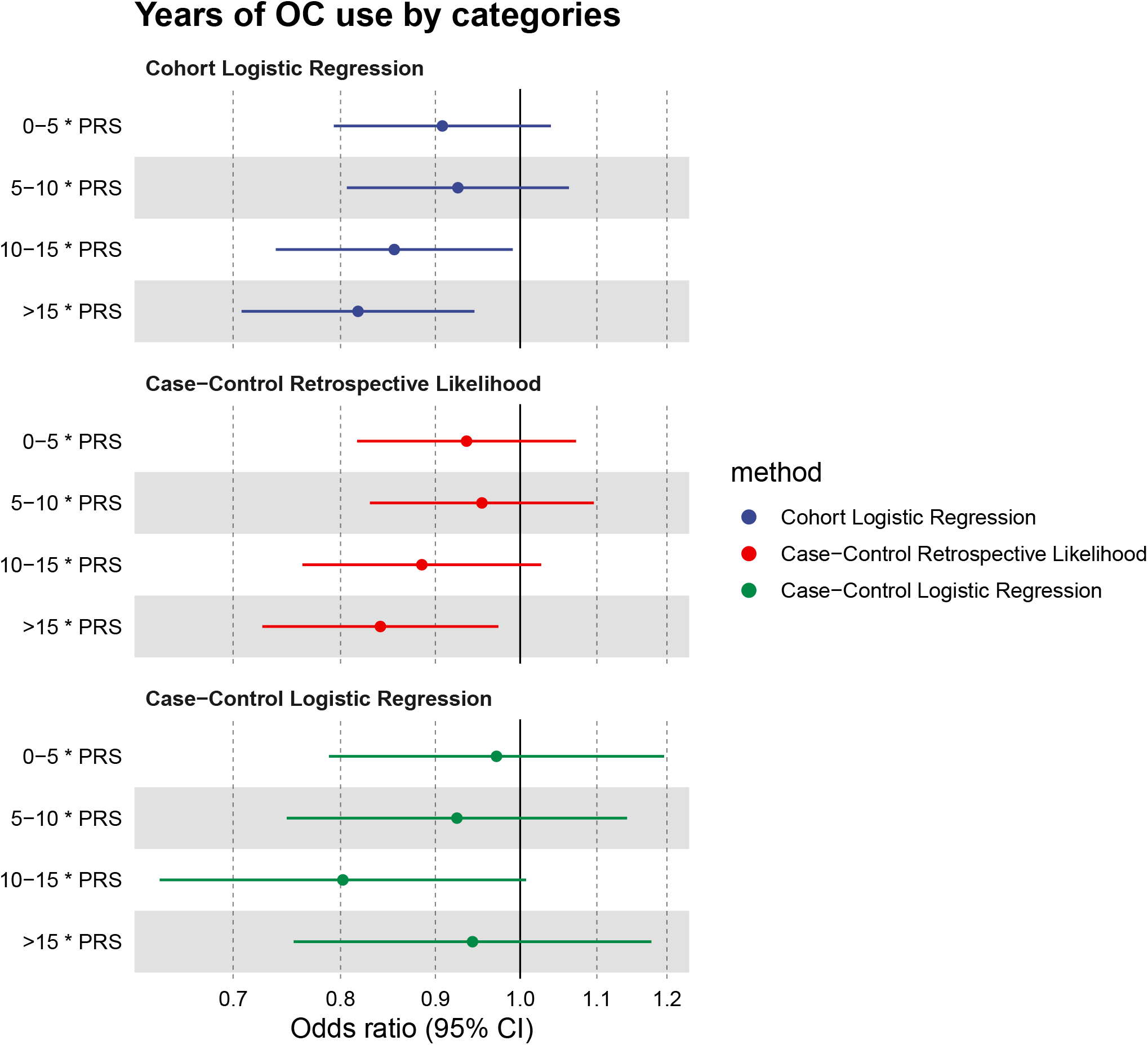
Odds ratio and 95% confidence interval for the categorized years of oral contraceptive use (YOC) in the risk model in the UK Biobank breast cancer data for postmenopausal women. Years of OC use are grouped into 5 categories: never, 0-5 years, 5-10 years, 10-15 years, and *>* 15 years. The model setting is the same as the continuous years of OC use model for all methods. The retrospective likelihood method and cohort logistic regression both showed a dose-response effect for YOC * PRS, with similar SD. CI, confidence interval.

We further observed that the retrospective-likelihood method detected the interaction between PRS and the birth location variable (birth.x: East coordinates) which were observed in the full cohort analysis (Table 2). For this term, the retrospective-likelihood produced a very similar result as the standard logistic regression analysis of case-control data. This result was expected as in the underlying PRS model we allowed for its correlation with this birth location variable. This was in contrast to the OC use, where our model assumed independence of PRS and OC use (conditional on stratification variables). Thus, the example illustrates the flexibility of the retrospective-likelihood method for trading off bias and efficiency in the estimation of interaction parameters by controlling the underlying gene-environment correlation parameters.

Finally, an inspection of the parameters of the PRS model indicated a significant correlation between PRS and the 4th PC, indicating the importance of incorporating population structures in modeling G-E correlation. The results on the complete sets of parameters from the disease model and the estimates of regression coefficients ***η***_***s***_ and 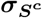 in the PRS model are presented in Appendix C Tables 5-8.

### 3.3 Analysis of UK Biobank Incident Colorectal Cancer

Table 3 shows results for colorectal cancer risk associated with PRS, age-groups and their interactions. The results on full sets of parameters can be found in Appendix C Tables 9 and 10. The logistic regression of the cohort data indicated statistically significant interactions between PRS and the age group 70+. We observed very similar patterns of operating characteristics of the different methods as in the breast cancer example. Both the retrospective-likelihood and the case-only method detected the PRS by age interactions, but standard logistic analysis of the case-control data failed to detect this pattern. For the estimation of the PRS main effect, the case-only method again produced an attenuated estimate and larger standard error compared to the retrospective-likelihood. We noticed a strong increasing dose-response relationship of age groups on the risk of colorectal cancer and a decreasing dose-response effect for the interaction terms of age with PRS for individuals older than 55.

**Table 3:**
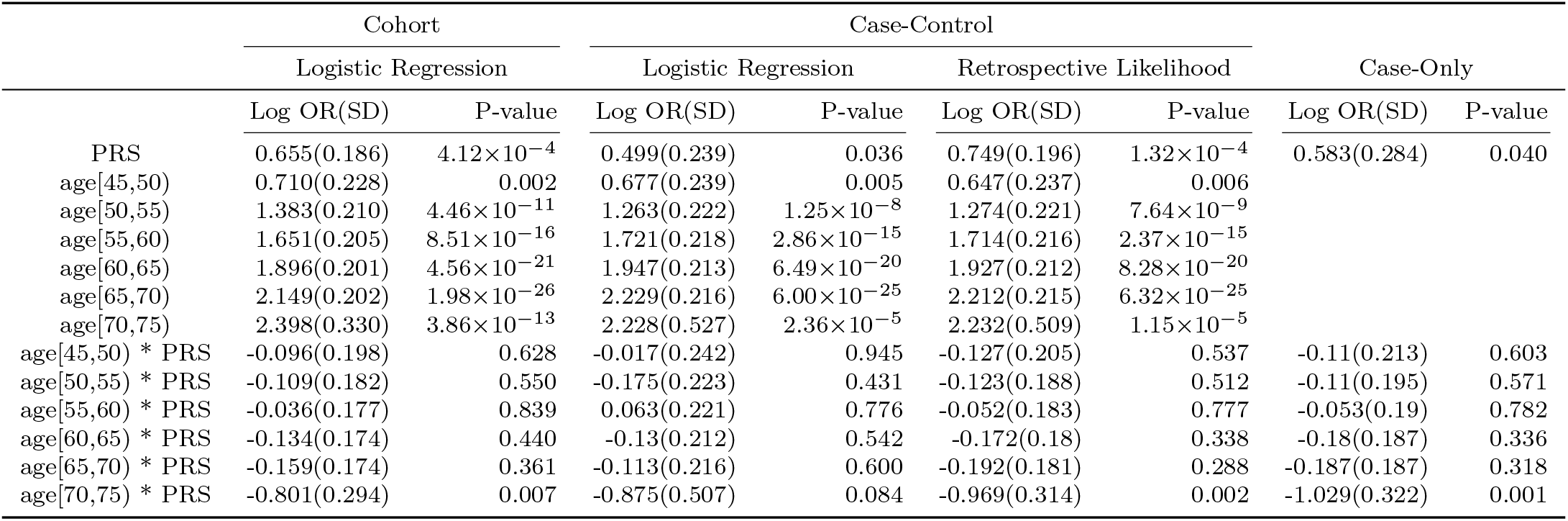
Parameter estimates of the log odds ratio (log OR), standard deviation, and p-values for the risk model in the UK Biobank colorectal cancer data. The reference group for age is [40,45). PRS-associated risks are presented in terms of per-SD unit. PRS, colorectal cancer-related polygenic risk score constructed from 95 SNPs.

## 4 Discussion

We have developed a novel method based on the semiparametric retrospective likelihood approach to analyze polygenic gene-environment interactions and correlations in case-control studies. The method is easy to implement and numerically stable that can incorporate large sets of covariates into both types of models. The method can flexibly exploit conditional independence assumption between PRS and E, adjusting for stratification factors and thus can effectively trade-off bias and efficiency for the estimation of different types of interaction parameters in the model. Conversely, the method can also produce efficient estimates of gene-environment correlation parameters by flexibly pooling data across case and controls according to specification of the underlying disease-risk model.

Our simulation studies indicate that the proposed method can substantially gain efficiency compared to standard logistic regression analysis of case-control studies by exploiting the gene-environment independence assumption. Yet the method can control for bias due to gene-environment correlation appearing due to mediation by complex stratification factors. Compared to two alternative methods for retrospective analysis, namely the case-only (Meisner et al., 2019) and the nonparametric regression approach (Stalder et al., 2017), the proposed method provides more flexibility to account for a complex set of stratification factors. Simulation studies also show that the proposed method can provide a more efficient analysis of gene-environment correlations compared to the traditional approach of using data only from controls. Our data analysis results are largely consistent with patterns observed in simulated data and further indicate the ability of the proposed method to reproduce results from cohort studies underlying case-control samples.

Our analysis of breast cancer risk in the UK Biobank reveals the potential presence of interactions between long-term OC use and polygenic risk for both the postmenopausal and premenopausal cohorts. Among premenopausal women, where OC use is positively associated with the risk of breast cancer, the pattern of interaction indicates that the risk associated with longer OC use is lower for women with higher polygenic risk and vice versa. In contrast, for postmenopausal women, the patterns of interaction and main effects indicate a protective effect of years of OC use among women at high polygenic risk. While interactions between OC use and the high-risk but rare BRCA1/2 mutations have been observed in the past (Lee et al., 2008), little has been known about such interactions in the context of polygenic risk associated with common variants. Our results require further confirmation in future studies.

Users need to be mindful of certain challenges and limitations associated with the proposed method. Our approach allows for gene-environment correlation through the incorporation of various stratification variables in the PRS model. It is, however, possible that there could be direct G-E associations for factors like BMI which have heritable components themselves. In the proposed method, the possibility of such correlations could be addressed by incorporating these variables in the mean model for the PRS, but then it is expected that the efficiency advantage of the proposed method over logistic regression will be lost. An alternative solution that could allow for direct G-E correlation and yet can retain the efficiency advantage of the retrospective likelihood method will be the following - adjust for an externally available polygenic score (PGS) for the environmental factor, but not the environmental factor itself, in the mean model for the PRS (Meisner et al., 2019). Under the assumption that the PRS of the disease is independent of E conditional on PGS and other stratification factors, this analysis is expected to provide unbiased and efficient estimates of gene-environment interaction parameters in the underlying disease-risk model.

Another limitation of the proposed method is the underlying rare disease assumption for simplifying likelihood calculations. For the analysis of single-SNP data, we (Chatterjee and Carroll, 2005) have previously shown that in the absence of a rare disease assumption, maximum-likelihood estimation can become numerically unstable unless external information on the marginal probability of the disease *π*_1_ is provided. In our setting, the rare disease assumption allows the derivation of the profile likelihood in an analytic form, taking advantage of the normality assumption for the distribution of PRS. It is, however, possible to implement the method without the rare disease assumption by evaluating the profile likelihood through numerical integration. However, external information on *π*_1_ should be provided to resolve any numerical issues associated with the estimation of the intercept parameter *α*, which is weakly identifiable from the profile likelihood under the gene-environment independence assumption.

In summary, we propose a highly flexible and efficient approach to modeling gene-environment interactions and correlations using PRS in case-control studies. The tool will aid better understanding of the interplay of PRS with various environmental and stratification factors, and thus ultimately could improve the interpretability and applicability of PRS for a variety of clinical and research applications.

## Acknowledgements

The research of Wang and Chatterjee was supported by the NIH grant 1R01HG010480. Additionally, research of Chatterjee was supported by NIH grants U01 CA249866, U01HG011719, 1U24OD023382-01.

## Code Availability

The R function and user’s guide of the proposed retrospective likelihood method, the computation codes of the simulation studies, and the codes and detailed results of the data applications to UK Biobank are available in https://github.com/ziqiaow/RetroGE

