## Supplementary Materials for "Joint Modeling of Gene-Environment Correlations and Interactions using Polygenic Risk Scores in Case-Control Studies"

### 1 Appendix A: Derivation of the Profile Likelihood and the Log-Likelihood Score Function

The profile likelihood can be derived as

$$L_i^R = f_{Z|S}(Z_i|\boldsymbol{S_i}) \frac{\exp(d_i\{\kappa + m(Z_i, \boldsymbol{E_i}, \boldsymbol{S_i}; \boldsymbol{\beta})\})/\tau(Z_i, \boldsymbol{E_i}, \boldsymbol{S_i}; \alpha, \boldsymbol{\beta})}{\sum_{d=0}^{1} \int f_{Z|S}(z'|\boldsymbol{S_i}) \exp(d\{\kappa + m(z', \boldsymbol{E_i}, \boldsymbol{S_i}; \boldsymbol{\beta})\})/\tau(z', \boldsymbol{E_i}, \boldsymbol{S_i}; \alpha, \boldsymbol{\beta}) dz'},$$

where $\kappa = \alpha + \log(N_1/N_0) - \log(\pi_1/\pi_0)$ and $\tau(Z, \boldsymbol{E}, \boldsymbol{S}; \alpha, \boldsymbol{\beta}) = 1 + \exp(\alpha + m(Z, \boldsymbol{E}, \boldsymbol{S}; \boldsymbol{\beta}))$.

Under rare disease assumption, the profile likelihood is simplified as

$$\begin{aligned}
L_i^R &= f_{Z|S}(Z_i|\boldsymbol{S_i}) \frac{\exp(D_i\{\kappa + m(Z_i, \boldsymbol{E_i}, \boldsymbol{S_i}; \boldsymbol{\beta})\})}{\sum_{d=0}^{1} \int f_{Z|S}(z'|\boldsymbol{S_i}) \exp(d\{\kappa + m(z', \boldsymbol{E_i}, \boldsymbol{S_i}; \boldsymbol{\beta})\}) dz'} \\
&= f_{Z|S}(Z_i|\boldsymbol{S_i}) \frac{\exp(D_i\{\kappa + m(Z_i, \boldsymbol{E_i}, \boldsymbol{S_i}; \boldsymbol{\beta})\})}{1 + \int \exp(\{\kappa + \beta_Z z' + \boldsymbol{\beta_E E_i} + \boldsymbol{\beta_S S_i} + \boldsymbol{\beta_{EZ} E_i} z' + \boldsymbol{\beta_{SZ} S_i} z'\}) \frac{1}{\sqrt{2\pi\sigma^2_{s^c,i}}} \exp\{-\frac{(z' - \boldsymbol{\eta_s}^T \boldsymbol{S_i})^2}{2\sigma^2_{s^c,i}}\} dz'} \\
&= f_{Z|S}(Z_i|\boldsymbol{S_i}) \frac{\exp(D_i\{\kappa + m(Z_i, \boldsymbol{E_i}, \boldsymbol{S_i}; \boldsymbol{\beta})\})}{1 + \exp\{\sigma^2_{s^c,i}(\beta_Z + \boldsymbol{\beta_{EZ} E_i} + \boldsymbol{\beta_{SZ} S_i})^2/2 + \eta_s^T s(\beta_Z + \beta_{EZ} e + \beta_{SZ} s) + \kappa + \boldsymbol{\beta_E E_i} + \boldsymbol{\beta_S S_i}\}} \\
&= \frac{\frac{1}{\sqrt{2\pi\sigma^2_{s^c,i}}} \exp\{-\frac{(Z_i - \boldsymbol{\eta_s}^T \boldsymbol{S_i})^2}{2\sigma^2_{s^c,i}} + D_i\{\kappa + m(Z_i, \boldsymbol{E_i}, \boldsymbol{S_i}; \boldsymbol{\beta})\}\}}{1 + \exp\{\sigma^2_{s^c,i}(\beta_Z + \boldsymbol{\beta_{EZ} E_i} + \boldsymbol{\beta_{SZ} S_i})^2/2 + \boldsymbol{\eta_s}^T \boldsymbol{S_i}(\beta_Z + \boldsymbol{\beta_{EZ} E_i} + \boldsymbol{\beta_{SZ} S_i}) + \kappa + \boldsymbol{\beta_E E_i} + \boldsymbol{\beta_S S_i}\}} \\
&= \frac{\frac{1}{\sqrt{2\pi\sigma^2_{s^c,i}}} \exp\{-\frac{(Z_i - \boldsymbol{\eta_s}^T \boldsymbol{S_i})^2}{2\sigma^2_{s^c}} + D_i\{\kappa + \beta_Z Z_i + \boldsymbol{\beta_E E_i} + \boldsymbol{\beta_S S_i} + \boldsymbol{\beta_{EZ} E_i} Z_i + \boldsymbol{\beta_{SZ} S_i} Z_i\}\}}{1 + \exp\{\sigma^2_{s^c,i}(\beta_Z + \boldsymbol{\beta_{EZ} E_i} + \boldsymbol{\beta_{SZ} S_i})^2/2 + \boldsymbol{\eta_s}^T \boldsymbol{S_i}(\beta_Z + \boldsymbol{\beta_{EZ} E_i} + \boldsymbol{\beta_{SZ} S_i}) + \kappa + \boldsymbol{\beta_E E_i} + \boldsymbol{\beta_S S_i}\}} \\
&= \frac{\frac{1}{\sqrt{2\pi\sigma^2_{s^c,i}}} \exp\{-\frac{(Z_i - \boldsymbol{\eta_s}^T \boldsymbol{S_i})^2}{2\sigma^2_{s^c,i}} + D_i\{R_1(\boldsymbol{E_i}, \boldsymbol{S_i}; \kappa, \boldsymbol{\beta_E}, \boldsymbol{\beta_S}) + Z_i R_2(\boldsymbol{E_i}, \boldsymbol{S_i}; \beta_Z, \boldsymbol{\beta_{EZ}}, \boldsymbol{\beta_{SZ}})\}\}}{1 + \exp\{\sigma^2_{s^c}\{R_2(\boldsymbol{E_i}, \boldsymbol{S_i}; \beta_Z, \boldsymbol{\beta_{EZ}}, \boldsymbol{\beta_{SZ}})\}^2/2 + \boldsymbol{\eta_s}^T \boldsymbol{S_i} R_2(\boldsymbol{E_i}, \boldsymbol{S_i}; \beta_Z, \boldsymbol{\beta_{EZ}}, \boldsymbol{\beta_{SZ}}) + R_1(\boldsymbol{E_i}, \boldsymbol{S_i}; \kappa, \boldsymbol{\beta_E}, \boldsymbol{\beta_S})\}},
\end{aligned}$$

where we define

$$\begin{aligned}
R_1(\boldsymbol{E}, \boldsymbol{S}; \kappa, \boldsymbol{\beta_E}, \boldsymbol{\beta_S}) &= \kappa + \boldsymbol{\beta_E E} + \boldsymbol{\beta_S S}; \\
R_2(\boldsymbol{E}, \boldsymbol{S}; \beta_Z, \boldsymbol{\beta_{EZ}}, \boldsymbol{\beta_{SZ}}) &= \beta_Z + \boldsymbol{\beta_{EZ} E} + \boldsymbol{\beta_{SZ} S}.
\end{aligned}$$

This model can further include the main effect of $S^c$ and interaction term between $Z$ and $S^c$ by formulating $R_1(\boldsymbol{E}, \boldsymbol{S}; \kappa, \boldsymbol{\beta_E}, \boldsymbol{\beta_S}, \beta_{S^c}) = \kappa + \boldsymbol{\beta_E E} + \boldsymbol{\beta_S S} + \beta_{S^c} S^c$ and $R_2(\boldsymbol{E}, \boldsymbol{S}; \beta_Z, \boldsymbol{\beta_{EZ}}, \boldsymbol{\beta_{SZ}}, \beta_{S^c Z}) = \beta_Z + \boldsymbol{\beta_{EZ} E} + \boldsymbol{\beta_{SZ} S} + \beta_{S^c Z} S^c$. Both $R_1(\cdot)$ and $R_2(\cdot)$ can remain flexible.

To estimate the unknown parameters defined as $\Omega_1 = (\kappa, \boldsymbol{\beta}^T)^T$ and $\Omega_2 = (\boldsymbol{\eta_s}^T, \boldsymbol{\sigma}^T_{\boldsymbol{S^c}})^T$, we use maximum likelihood estimation with the input of score function. The log-likelihood is

$$\log L^R = \sum_{i=1}^{N} \left\{ \log f_{Z|S}(Z_i|\boldsymbol{S_i}) + D_i\{\kappa + m(Z_i, \boldsymbol{E_i}, \boldsymbol{S_i}; \boldsymbol{\beta})\} - \log \sum_{d=0}^{1} \int f_{Z|S}(z'|\boldsymbol{S_i}) \exp(d\{\kappa + m(z', \boldsymbol{E_i}, \boldsymbol{S_i}; \boldsymbol{\beta})\}) dz' \right\}.$$

Define

$$f_1(D_i, Z_i, \boldsymbol{E_i}, \boldsymbol{S_i}; \Omega_1) = D_i\{\kappa + m(Z_i, \boldsymbol{E_i}, \boldsymbol{S_i}; \boldsymbol{\beta})\};$$
$$f_2(\boldsymbol{E_i}, \boldsymbol{S_i}; \Omega_1) = \sum_{d=0}^{1} \int f_{Z|S}(z'|\boldsymbol{S_i}) \exp(d\{\kappa + m(z', \boldsymbol{E_i}, \boldsymbol{S_i}; \boldsymbol{\beta})\})dz'.$$

The score function for the log-likelihood can be expressed as

$$\frac{\partial \log L^R}{\partial \Omega} = \sum_{i=1}^{N} \left\{ \frac{\partial \log f_{Z|S}(z_i|\boldsymbol{S_i})}{\partial \Omega_2} + \frac{\partial f_1(D_i, Z_i, \boldsymbol{E_i}, \boldsymbol{S_i}; \Omega_1)}{\partial \Omega_1} - \frac{\partial f_2(\boldsymbol{E_i}, \boldsymbol{S_i}; \Omega_1)/\partial \Omega_1}{f_2(\boldsymbol{E_i}, \boldsymbol{S_i}; \Omega_1)} \right\},$$

The derivative can be simply calculated, for each parameter we have

$$\frac{\partial \log L^R}{\partial \kappa} = \sum_{i=1}^{N} \{D_i - (1 - \frac{1}{f_2})\},$$
$$\frac{\partial \log L^R}{\partial \beta_Z} = \sum_{i=1}^{N} \left\{ D_i Z_i - (1 - \frac{1}{f_2})\{\sigma^2_{S^c,i} R_2(\boldsymbol{E_i}, \boldsymbol{S_i}; \beta_Z, \boldsymbol{\beta_{EZ}}, \boldsymbol{\beta_{SZ}}) + \boldsymbol{\eta_s}^T \boldsymbol{S_i}\} \right\},$$
$$\frac{\partial \log L^R}{\partial \boldsymbol{\beta_E}} = \sum_{i=1}^{N} \{D_i \boldsymbol{E_i} - (1 - \frac{1}{f_2})\boldsymbol{E_i}\},$$
$$\frac{\partial \log L^R}{\partial \boldsymbol{\beta_S}} = \sum_{i=1}^{N} \{D_i \boldsymbol{S_i} - (1 - \frac{1}{f_2})\boldsymbol{S_i}\},$$
$$\frac{\partial \log L^R}{\partial \boldsymbol{\beta_{EZ}}} = \sum_{i=1}^{N} \left\{ D_i \boldsymbol{E_i} Z_i - (1 - \frac{1}{f_2})\{\sigma^2_{S^c,i} R_2(\boldsymbol{E_i}, \boldsymbol{S_i}; \beta_Z, \boldsymbol{\beta_{EZ}}, \boldsymbol{\beta_{SZ}})\boldsymbol{E_i} + \boldsymbol{\eta_s}^T \boldsymbol{S_i}\boldsymbol{E_i}\} \right\},$$
$$\frac{\partial \log L^R}{\partial \boldsymbol{\beta_{SZ}}} = \sum_{i=1}^{N} \left\{ D_i \boldsymbol{S_i} Z_i - (1 - \frac{1}{f_2})\{\sigma^2_{S^c,i} R_2(\boldsymbol{E_i}, \boldsymbol{S_i}; \beta_Z, \boldsymbol{\beta_{EZ}}, \boldsymbol{\beta_{SZ}})\boldsymbol{S_i} + \boldsymbol{\eta_s}^T \boldsymbol{S_i}\boldsymbol{S_i}\} \right\},$$
$$\frac{\partial \log L^R}{\partial \boldsymbol{\eta_s}} = \sum_{i=1}^{N} \left\{ \frac{(Z_i - \boldsymbol{\eta_s}^T \boldsymbol{S_i})\boldsymbol{S_i}}{\boldsymbol{\sigma^2_{S^c,i}}} - (1 - \frac{1}{f_2})\{R_2(\boldsymbol{E_i}, \boldsymbol{S_i}; \beta_Z, \boldsymbol{\beta_{EZ}}, \boldsymbol{\beta_{SZ}})\boldsymbol{S_i}\} \right\},$$
$$\frac{\partial \log L^R}{\partial \boldsymbol{\sigma_{S^c}}} = \sum_{i=1}^{N} \boldsymbol{\sigma_{S^c}} \left\{ (Z_i - \boldsymbol{\eta_s}^T \boldsymbol{S_i})^2 \boldsymbol{\sigma_{S^c}^{-3}} - \frac{1}{\boldsymbol{\sigma_{S^c}}} - (1 - \frac{1}{f_2})\{R_2^2(\boldsymbol{E_i}, \boldsymbol{S_i}; \beta_Z, \boldsymbol{\beta_{EZ}}, \boldsymbol{\beta_{SZ}})\} \right\}.$$

### 2 Appendix B: Simulation Results for All Parameters in the Model

#### 2.1 Parameter Values and Detailed Descriptions

We first simulated a cohort of $10^6$ individuals with a population disease prevalence of $\pi = \Pr(D = 1) = 0.01$ using a disease model assuming a logistic regression model of the form:

$$\text{pr}(D = 1|Z, E, S) = \alpha + \beta_Z Z + \beta_{E_1} E_1 + \beta_{E_2} E_2 + \beta_{E_1 Z} E_1 Z + \beta_{E_2 Z} E_2 Z + \boldsymbol{\beta}_{\boldsymbol{S}}^T \boldsymbol{S}. \quad (1)$$

In scenario 1, we assumed a model without variables $S$ and we assumed that $Z$ is independent of $E$ with $Z \sim N(\mu, \sigma^2)$. We assessed the type I error with $\beta_{E_1 Z} = 0, \beta_{E_2 Z} = 0$. The other coefficients are $\beta_Z = 0.450, \beta_{E_1} = 0.15, \beta_{E_2} = -0.07$. These parameters were retrieved from our data analysis fitting logistic regression model on the cohort study of breast cancer in the UK Biobank. The odds ratio of PRS is around $\exp(0.45) = 1.57$. The intercept term is set as $\alpha = -5$ to make the population disease prevalence of approximately 0.01.

In scenario 2, we added $\boldsymbol{S}$ as covariates in the logistic regression model (1). We added one stratification factor $S_1$ of 3 levels and a continuous variable $S_2$ so that $Z \perp\!\!\!\perp E|(S_1, S_2)$. For the model, we create dummy variables for $S_1$ as $S_{11}, S_{12}$. Here, we let $m(Z, \boldsymbol{E}, \boldsymbol{S}; \beta) = \beta_Z Z + \beta_{E_1} E_1 + \beta_{E_2} E_2 + \beta_{E_1 Z} E_1 Z + \beta_{E_2 Z} E_2 Z + \beta_{S_{11}} S_{11} + \beta_{S_{12}} S_{12} + \beta_{S_2} S_2$, with PRS $Z|S \sim N(\eta_s^T S, \sigma^2(S_1))$, $E_1 \sim Bin(0.745)$, $E_2 \sim N(0, 1)$, $S_{11}$ and $S_{12}$ are two dummy variables for $S^c$ of 3 levels, and $S_2 \sim N(0, 1)$ is a continuous variable. The coefficients are $\beta_Z = 0.450, \beta_{E_1} = 0.15, \beta_{E_2} = -0.07, \beta_{E_1 Z} = 0, \beta_{E_2 Z} = -0.40, \beta_{S_{11}} = 0.1, \beta_{S_{12}} = 0.3, \beta_{S_2} = 0.5$. $\boldsymbol{\eta_s} = c(1, 0.2, 0.3, 0.2)$, 1 is the coefficient for the intercept term and $\boldsymbol{\sigma_{S_1}} = c(0.25, 0.5, 1)$ for each stratum in $S_1$.

Scenario 3 is similar to scenario 2, but adding the interaction terms between $S$ and $Z$ into the model, i.e., $m(Z, \boldsymbol{E}, \boldsymbol{S}; \beta) = \beta_Z Z + \beta_{E_1} E_1 + \beta_{E_2} E_2 + \beta_{E_1 Z} E_1 Z + \beta_{E_2 Z} E_2 Z + \beta_{S_{11}} S_{11} + \beta_{S_{12}} S_{12} + \beta_{S_2} S_2 + \beta_{S_{11} Z} S_{11} Z + \beta_{S_{12} Z} S_{12} Z + \beta_{S_2 Z} S_2 Z$. The interaction term coefficients are $\beta_{S_{11} Z} = 0, \beta_{S_{12} Z} = 0.1, \beta_{S_2 Z} = 0.2$. All the other parameters are the same as those in scenario 2.

Scenario 4 is a population logistic regression model without variables $S$ but we assumed that $Z|S \sim N(\boldsymbol{\eta_s}^T \boldsymbol{S}, \boldsymbol{\sigma^2_{S_1}})$ as described in scenario 2. In addition, $S_2$ and $E_2$ follow a bivariate normal distribution with a correlation of 0.4. Scenario 5 is similar to scenario 4 but adds $\boldsymbol{S}$ as covariates into the logistic regression model. In scenario 6, we eliminate the independence assumption between PRS and environmental variables by only including $\boldsymbol{S}$ into the population model so that PRS depends on all environmental variables. For scenario 6, we expect our model to perform similarly to the logistic regression model in a case-control setting as there is no independent covariate to PRS that we could exploit in our method.

#### 2.2 Type I Error Assessments

| Type I Error | Retro.Likelihood | Logistic.Regression | Nonparametrics | Case.Only.Population | Case.Only |
|---|---|---|---|---|---|
| Scenario 1: PRS x $E_1$ | 0.042 | 0.057 | 0.047 | 0.042 | 0.042 |
| Scenario 1: PRS x $E_2$ | 0.038 | 0.041 | 0.044 | 0.042 | 0.042 |
| Scenario 2: PRS x $E_1$ | 0.070 | 0.062 | 0.067 | 0.066 | 0.066 |
| Scenario 3: PRS x $E_1$ | 0.035 | 0.037 | 0.042 | 0.039 | 0.039 |
| Scenario 3: PRS x $S_{11}$ | 0.050 | 0.054 | 1.000 | 0.961 | 0.961 |
| Scenario 4: PRS x $E_1$ | 0.055 | 0.047 | 0.051 | 0.054 | 0.054 |
| Scenario 5: PRS x $E_1$ | 0.058 | 0.039 | 0.057 | 0.054 | 0.054 |
| Scenario 6: PRS x $S_{11}$ | 0.048 | 0.039 | 1.000 | 0.992 | 0.992 |

Table 1: Type I error rates at a nominal significance level of 0.05 for all 6 scenarios in the simulation study. Scenario 1 assumes PRS and $E_1, E_2$ are independent; scenario 2 assumes PRS $\perp\!\!\!\perp E|S$ and $S$ are included as covariates in the disease model; scenario 3 assumes PRS $\perp\!\!\!\perp E|S$ and $S$ are included as both covariates and PRS x $S$ interaction terms in the disease model; scenario 4 assumes PRS $\perp\!\!\!\perp E|S$ and $f(E, S)$ follows a bivariate normal distribution; scenario 5 assumes PRS $\perp\!\!\!\perp E|S$ and $f(E, S)$ follows a bivariate normal distribution and $S$ are included as both covariates and PRS x $S$ interaction terms in the disease model; scenario 6 assumes no independent environmental variable in the model, i.e., only $S$ and its interaction terms with PRS are included in the disease model.

#### 2.3 The Correlation Assessments between PRS and $S$ Using Weighted and Adaptive Weighted Methods

Supposed we denote the case-only and control-only linear regression coefficients for the parameter of interest to be $\hat{\beta}_{CA}$ and $\hat{\beta}_{CO}$. Then the weighted estimator is

$$\hat{\beta}_W = w_{cc}\hat{\beta}_{CO} + (1 - w_{cc})\hat{\beta}_{CA},$$

where $w_{cc} = \hat{\sigma}^2_{CA}/(\hat{\sigma}^2_{CA} + \hat{\sigma}^2_{CO})$. Here, $\hat{\sigma}^2_W = \hat{Var}(\hat{\beta}_W) = \hat{\sigma}^2_{CA}\hat{\sigma}^2_{CO}/(\hat{\sigma}^2_{CA} + \hat{\sigma}^2_{CO})$.

The adaptive weighted estimator for the correlation coefficient between PRS and $S$ is proposed as

$$\hat{\beta}_{AW} = \frac{\hat{\beta}^2_{SZ}}{\hat{\sigma}^2_{CO} + \hat{\beta}^2_{SZ}}\hat{\beta}_{CO} + \frac{\hat{\sigma}^2_{CO}}{\hat{\sigma}^2_{CO} + \hat{\beta}^2_{SZ}}\hat{\beta}_W,$$

where $\hat{\beta}^2_{SZ}$ is the estimated interaction term between PRS and $S$ in the disease model using standard logistic regression. Supposed we neglect the variability of $\hat{\beta}^2_{SZ}$, $\hat{\sigma}^2_{CO}$, and $\hat{\sigma}^2_{CA}$, we obtain the following variance estimate,

$$
\begin{aligned}
\hat{\sigma}^2_{AW} &= \hat{Var}(\hat{\beta}_{AW}) \\
&= \hat{Var}\left(\frac{\hat{\beta}^2_{SZ}}{\hat{\sigma}^2_{CO} + \hat{\beta}^2_{SZ}}\hat{\beta}_{CO} + \frac{\hat{\sigma}^2_{CO}}{\hat{\sigma}^2_{CO} + \hat{\beta}^2_{SZ}}(w_{cc}\hat{\beta}_{CO} + (1 - w_{cc})\hat{\beta}_{CA})\right) \\
&= \left(\frac{\hat{\beta}^2_{SZ} + \hat{\sigma}^2_{CO}w_{cc}}{\hat{\sigma}^2_{CO} + \hat{\beta}^2_{SZ}}\right)^2 \hat{\sigma}^2_{CO} + \left(\frac{\hat{\sigma}^2_{CO}(1 - w_{cc})}{\hat{\sigma}^2_{CO} + \hat{\beta}^2_{SZ}}\right)^2 \hat{\sigma}^2_{CA}.
\end{aligned}
$$

Note that as $\hat{\beta}^2_{SZ} \to 0$, $\hat{\beta}_{AW} \to \hat{\beta}_W$, and as $\hat{\beta}^2_{SZ} \to \infty$, $\hat{\beta}_{AW} \to \hat{\beta}_{CO}$. The theoretical background and rationale of the proposed methods can be found in Li et al. (2010).

#### 2.4 Estimation of Sigma in PRS Model

| | $\sigma_{S_1} Strata1$ | | | $\sigma_{S_1} Strata2$ | | | $\sigma_{S_1} Strata3$ | | |
|---|---|---|---|---|---|---|---|---|---|
| | Bias | SD | Empirical SD | Bias | SD | Empirical SD | Bias | SD | Empirical SD |
| Scenario2 | -0.001 | 0.010 | 0.010 | -0.002 | 0.019 | 0.020 | -0.010 | 0.034 | 0.035 |
| Scenario3 | -0.001 | 0.011 | 0.011 | -0.003 | 0.020 | 0.019 | -0.025 | 0.033 | 0.034 |
| Scenario4 | -0.001 | 0.010 | 0.010 | -0.002 | 0.019 | 0.020 | -0.011 | 0.035 | 0.035 |
| Scenario5 | -0.001 | 0.010 | 0.010 | -0.001 | 0.019 | 0.019 | -0.012 | 0.035 | 0.034 |
| Scenario6 | -0.001 | 0.010 | 0.010 | -0.002 | 0.020 | 0.019 | -0.011 | 0.034 | 0.035 |

Table 2: Performance of retrospective likelihood method for the variance estimates in the PRS model in simulation studies. Bias, standard deviation (SD) and empirical SD for $\sigma_{S_1}$ in the simulation scenarios 2-6 where we assume PRS follows $Z|\boldsymbol{S} \sim N(\boldsymbol{\eta}_{\boldsymbol{s}}^T\boldsymbol{S}, \boldsymbol{\sigma}^{\mathbf{2}}_{\boldsymbol{S_1}})$. $S_1$ contains three strata with true value of $\boldsymbol{\sigma}_{\boldsymbol{S_1}} = c(0.25, 0.5, 1)$ for each stratum.

#### 2.5 MSE of the Main Effect and Interaction Parameters between PRS and Environmental Variables

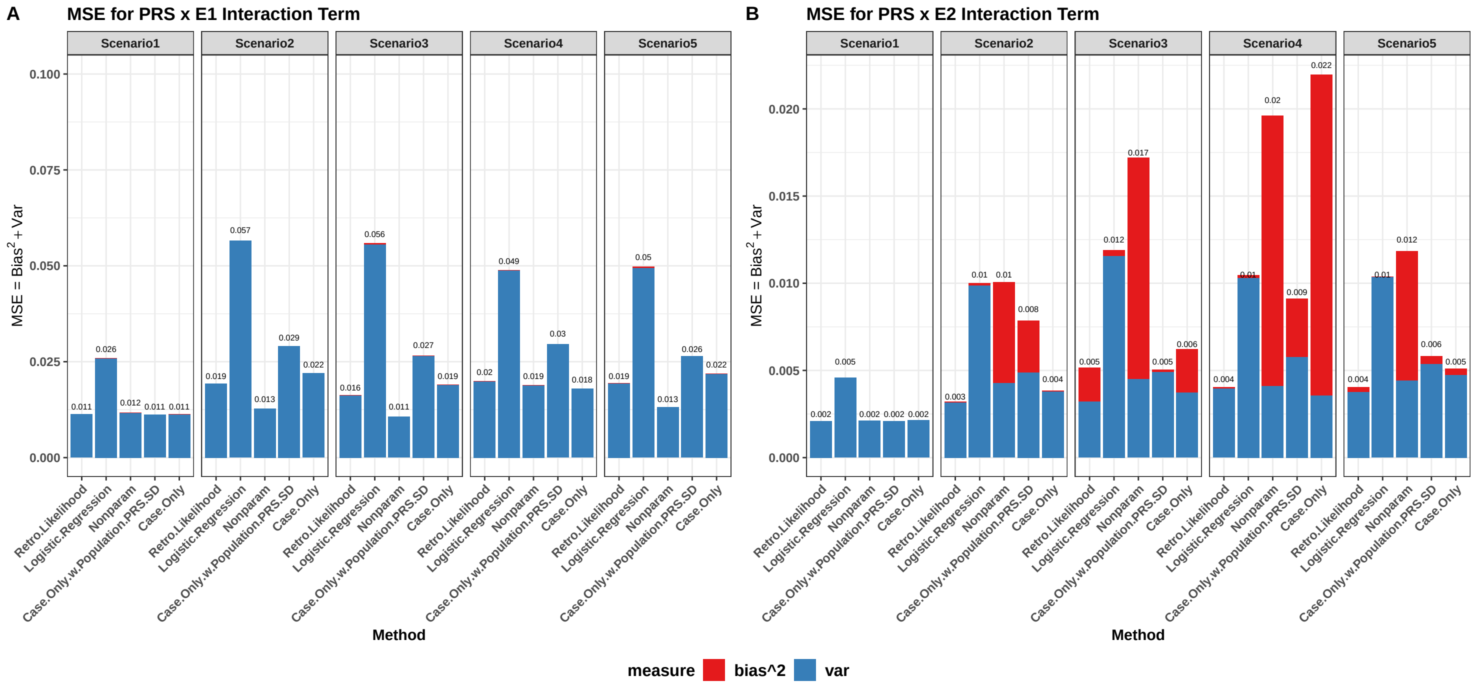

Figure 1: Mean squared error (MSE) of the interaction parameters between PRS and environmental variables $\boldsymbol{E}$ in the disease model in simulation scenarios 1-5.

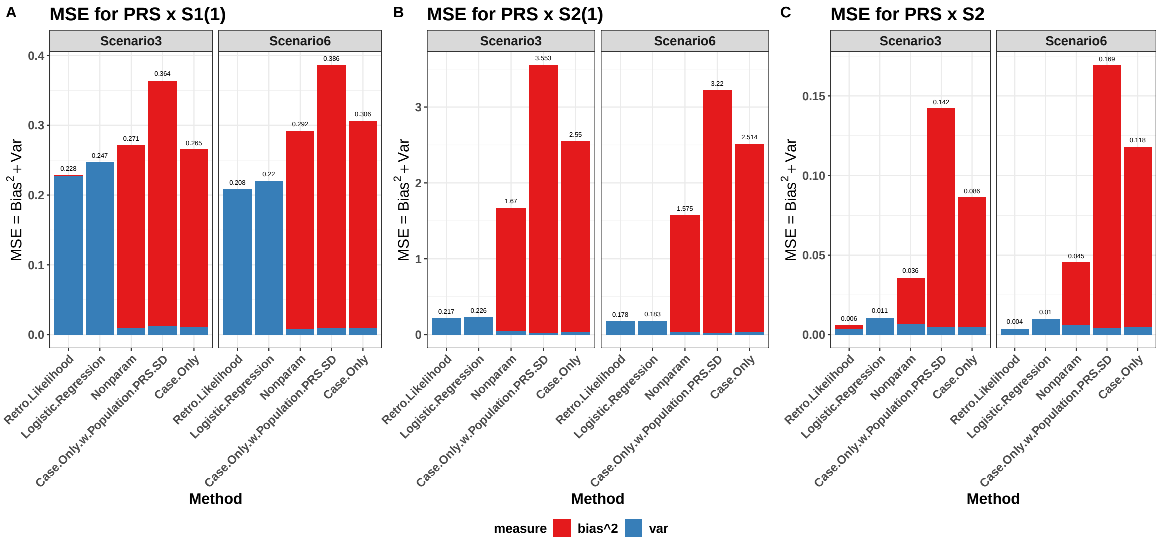

Figure 2: Mean squared error (MSE) of the interaction parameters between PRS and stratification variables $\boldsymbol{S}$ in the disease model in simulation scenarios 3 and 6. $S_{1(1)}$ and $S_{2(1)}$ denote the two dummy variables of $S_1$ in the mean function of PRS model.

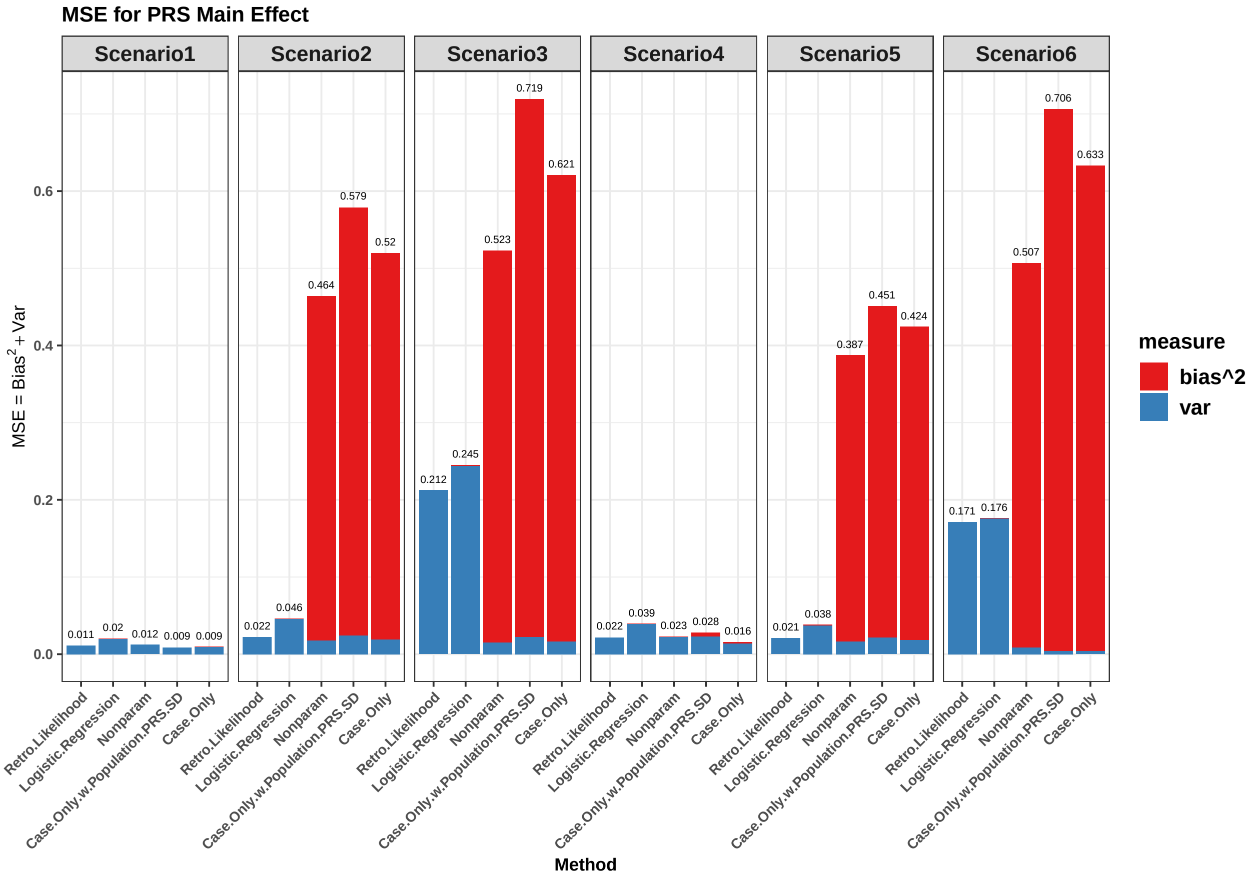

Figure 3: Mean squared error (MSE) of the main effect parameter of PRS in the disease model in simulation scenarios 1-6.

##### 2.6 Estimation of Main Effect Parameters for Environmental Variables

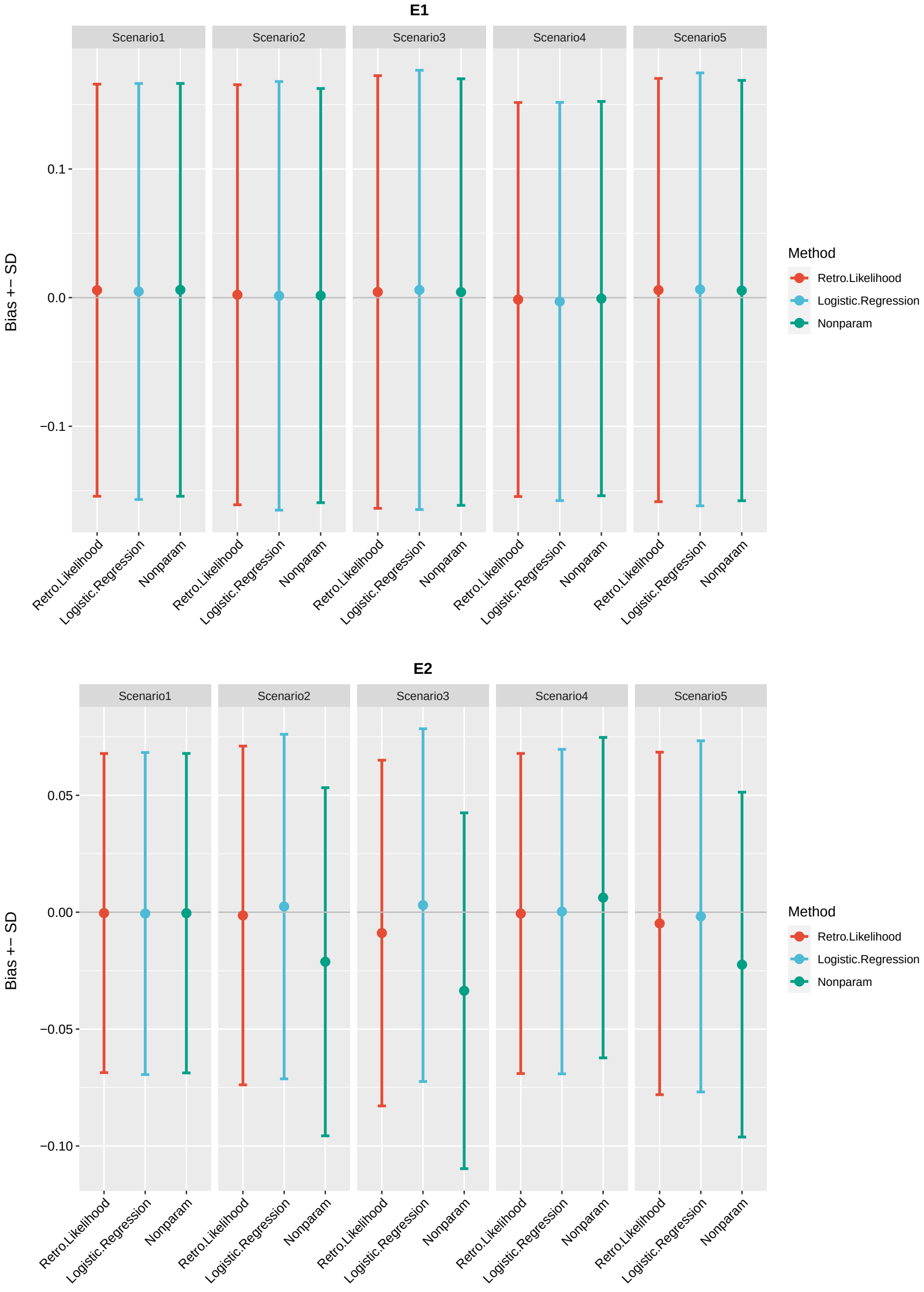

Figure 4: Performance of alternative methods for the main effect parameter estimates of environmental variables $\boldsymbol{E}$ in the disease model in simulation studies. The bias and estimated standard deviation (SD) of the main effect parameter $\beta_{E1}, \beta_{E2}$ in simulation scenarios 1-5. Note that the case-only method cannot estimate the parameters for the main effect of the environmental variables. In the figure, each dot represents bias and the error bar is SD.

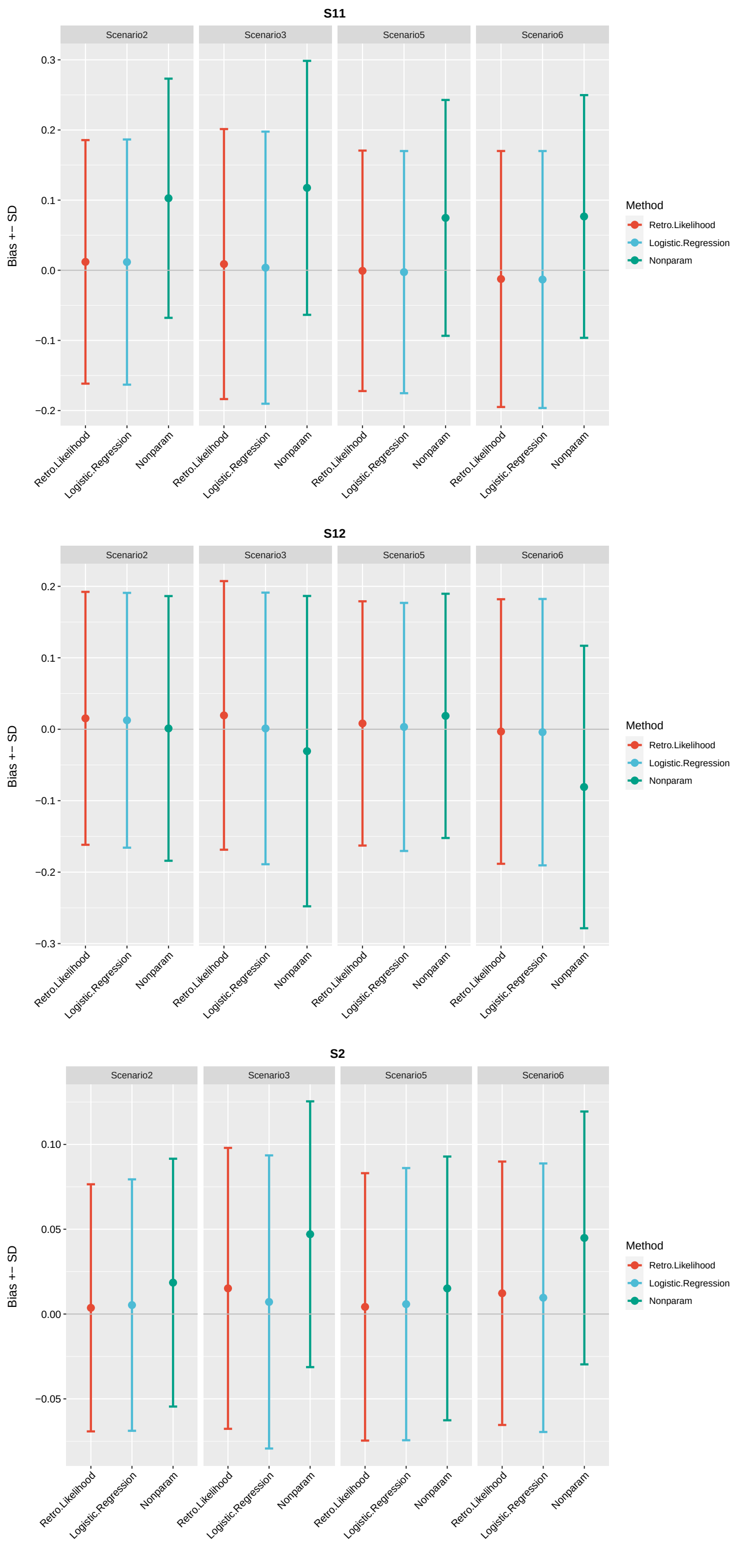

Figure 5: Performance of alternative methods for the main effect parameter estimates of environmental variables $\boldsymbol{S}$ in the disease model in simulation studies. The bias and estimated standard deviation (SD) of the main effect parameter $\beta_{S_{11}}, \beta_{S_{12}}, \beta_{S_2}$ in simulation scenarios 2,3,5,6. Note that the case-only method cannot estimate the parameters for the main effect of the environmental variables. In the figure, each dot represents bias and the error bar is SD.

### 3 Appendix C: Estimated Parameters in UK Biobank Data Analysis

#### 3.1 Details of the Data Analysis for Incident Breast Cancer

The main goal here is to discover PRS x $E$ interactions on the risk of breast cancer among premenopausal and postmenopausal women, respectively. The PRS is constructed using imputed genotype data based on the weighted sum of summary statistics of 313 SNPs reported in an external study Mavaddat et al. (2019) with tools provided by PGS Catalog in genome build GRCh37 (Polygenic Score (PGS) ID: PGS000004), see details in (Lambert et al., 2021; PGS Catalog Team, 2022). After removing ambiguous and duplicated SNPs, 276 SNPs were used for the PRS calculation. The odds ratio (OR) per standard deviation (SD) change of PRS for incident breast cancer in UK Biobank is 1.58 with 95% confidence interval (CI) (1.53,1.62) for all women after adjusting for the top 10 PCs; 1.56 (1.50,1.62) for postmenopausal women and 1.63 (1.52,1.75) for premenopausal women. These results are close to the OR 1.61 (1.57, 1.65) per SD change reported by Mavaddat et al. (2019).

In the disease model, we considered interactions between PRS and 10 previously reproduced classical risk factors (Kapoor et al., 2020; Pal Choudhury et al., 2019) including age (field 21022 in UK Biobank data), parity (field 2734), age at first birth (fields 2754 and 3872), alcohol intake (field 1558), height (field 12144), body mass index (BMI, field 21001), age at menarche (field 2714), years of OC use, age at menopause (for postmenopausal women, field 3581), and hormone replacement therapy (HRT) use (for postmenopausal women, field 2814). Years of OC use are defined by taking the difference of age when last used OC (field 2804) and age started OC (field 2794). We additionally included interactions of PRS and geographic information, the North and East coordinates of individual's birth location (fields 129 and 130), recorded on metre grid scales treated as continuous variables. Apart from the main effect and interaction terms, we adjusted for the top 10 PCs (field 22009) and categorical assessment centres (field 54) as additional covariates in the model. We make the assumption of independence between PRS and the environmental exposures conditional on $S$ = (Top 10 PCs, North coordinates of birth location, East coordinates of birth location) and stratification factor $S^c$ = (Assessment Centres). Thus the PRS model is as follows:

$$Z|S \sim N(\eta_0 + \eta_{\text{birth.x}}\text{birth.x} + \eta_{\text{birth.y}}\text{birth.y} \\ + \eta_{pc_1}PC_1 + \cdots + \eta_{pc_{10}}PC_{10}, \sigma^2_{\text{Assessment Centres}}), \quad (2)$$

where birth.x and birth.y are the East and North coordinates of birth location respectively.

We selected unrelated white female individuals with consent from the UK Biobank study. To identify breast cancer patients, we used the International Classification of Diseases (ICD) for types of cancer in UK Biobank data: ICD9 (field 40013) and ICD10 (field 40006). These include ICD-10 codes C50.0-C50.9 and ICD-9 codes 1740-1749. Breast cancer in situ patients (ICD-10 codes D05.0, D05.1, D05.7, D05.9, and ICD-9 code 2330) were excluded from the analysis. For risk assessment in this work, we only included incident breast cancer cases of patients who were diagnosed post-recruitment by comparing the age at cancer diagnosis (field 40008) with the age at recruitment (field 21022).

This yielded 4 170 incident breast cancer cases and 193 511 controls in the study. Based on the data available from self-reported menopause status (field 2724), 2 662 breast cancer incidents and 116 965 controls were included for postmenopausal women; 852 breast cancer incidents and 45 964 controls were selected for premenopausal women. After removing individuals with missing data for the above-described risk factors in the model, there were 2 024 cases and 89 241 controls for the postmenopausal cohort; 704 cases and 37 785 controls for the premenopausal cohort for analysis. We simulated a case-control study with all 2 024 cases and 2 024 randomly sampled controls from the 89 241 controls for the postmenopausal cohort and similarly for the premenopausal cohort we randomly selected 704 controls into the case-control design.

We considered 2 designs: 1) The prospective cohort design as described; 2) a mock case-control study with all 2 024 cases and 2 024 randomly sampled controls from the 89 241 controls for postmenopausal cohort and similarly for the premenopausal cohort we randomly selected 704 controls into the case-control design. We did 5 sets of analyses for comparisons: 1) we analyzed the cohort study with standard logistic regression with the interaction terms of PRS and the 10 classical risk factors and geographical regions adjusting for PCs and assessment centres; 2) we applied the proposed retrospective likelihood method in the case-control design assuming the same model as the cohort design and the PRS model in main text formula (5); 3) we applied logistic regression in the case-control design with the same model as in the cohort design; and 4) case-only method in the case-control study for the same covariates.

#### 3.2 Details of the Data Analysis for Incident Colorectal Cancer

We investigated the PRS x $E$ effect on the risk of colorectal cancer. The PRS is constructed using the summary statistics of 95 SNPs (Polygenic Score (PGS) ID: PGS000765) from an external study (Huyghe et al., 2019). After removing ambiguous and duplicated SNPs, 86 SNPs were used for the PRS calculation. The OR per SD change of PRS after adjusting for the top 10 PCs is 1.51 (1.45,1.57) for incident colorectal cancer in UK Biobank. We modeled interactions between PRS and lifestyle-based risk factors reported by a prospective study (Aleksandrova et al., 2021), including age, gender, alcohol intake, height, waist circumference (field 48), smoking status (field 20116), red meat intake, vegetable intake, process meat intake per week (field 1349), and the activity levels, defined by International Physical

Activity Questionnaire (IPAQ) (field 22032). The activity levels are aggregated categorical values of low, moderate, and high levels summarized based on 4 types of activities across leisure time, domestic, work-related, and transport-related physical activities. For the red meat intake, we considered the total beef (field 1369), lamb (field 1379), and pork (field 1389) intake frequency per week, by summing up the ordinal variables and treating it as a continuous variable. The vegetable intake is the summation of cooked (field 1289) and raw vegetables (field 1299) intake quantities per day. We additionally included interactions of PRS and geographic birth location and adjusted for the top 10 PCs and assessment centres as additional covariates in the disease model. The conditional independence structures of environmental variables in the PRS model are defined as (2).

Colorectal cancer is defined by malignant neoplasm of the colon (ICD-10 codes C18.0–18.9; ICD-9 codes 1530-1539), in the rectosigmoid junction (ICD-10 code C19; ICD-9 code 1540), or in the rectum (ICD-10 code C20; ICD-9 code 1541) (Aleksandrova et al., 2021). In total there are 2 370 incident cases and 379 036 controls. We selected unrelated white individuals with consent from the UK Biobank study. After removing the individuals with incomplete data, we used 1 743 incident cases and 282 429 controls for the cohort study and randomly selected 1 743 controls for the case-control design. We performed the same sets of analyses as we did on the breast cancer data.

#### 3.3 Parameter Estimates of UK Biobank Breast Cancer and Colorectal Cancer Data

| | Cohort | | Case-Control | | | |
|---|---|---|---|---|---|---|
| Post-Menopause | Logistic Regression | | Logistic Regression | | Retrospective Likelihood | |
| | Estimate(SD) | P-value | Estimate(SD) | P-value | Estimate(SD) | P-value |
| oc 0-10 | -0.001(0.067) | 0.983 | -0.122(0.089) | 0.170 | -0.118(0.087) | 0.177 |
| oc >10 | 0.055(0.07) | 0.434 | -0.05(0.094) | 0.596 | -0.057(0.092) | 0.535 |
| oc 0-10 * PRS | -0.087(0.06) | 0.146 | -0.054(0.093) | 0.562 | -0.057(0.061) | 0.344 |
| oc >10 * PRS | -0.18(0.064) | 0.005 | -0.13(0.098) | 0.183 | -0.149(0.064) | 0.021 |
| | Estimate(SD) | P-value | Estimate(SD) | P-value | Estimate(SD) | P-value |
| oc 0-5 | -0.035(0.076) | 0.646 | -0.115(0.101) | 0.257 | -0.115(0.1) | 0.250 |
| oc 5-10 | 0.037(0.078) | 0.637 | -0.133(0.104) | 0.198 | -0.124(0.102) | 0.223 |
| oc 10-15 | 0.066(0.082) | 0.422 | 0.036(0.111) | 0.749 | 0.043(0.11) | 0.698 |
| oc >15 | 0.047(0.08) | 0.556 | -0.119(0.108) | 0.268 | -0.144(0.106) | 0.172 |
| oc 0-5 * PRS | -0.097(0.069) | 0.160 | -0.029(0.106) | 0.782 | -0.067(0.069) | 0.337 |
| oc 5-10 * PRS | -0.077(0.07) | 0.272 | -0.079(0.108) | 0.466 | -0.048(0.071) | 0.503 |
| oc 10-15 * PRS | -0.156(0.075) | 0.037 | -0.22(0.116) | 0.058 | -0.122(0.076) | 0.106 |
| oc >15 * PRS | -0.201(0.074) | 0.006 | -0.059(0.113) | 0.602 | -0.174(0.075) | 0.020 |

Table 3: Parameter estimates of the log odds ratio, standard deviation, and p-values for the disease-risk model in the UK Biobank breast cancer data for postmenopausal women. Years of OC use is categorized into two sets: 1) never, 0-10 years, and $>$ 10 years. 2) never, 0-5 years, 5-10 years, 10-15 years, and $>$ 15 years. The reference group is women who have never taken OC. The model setting is the same as the continuous years of OC use model for all methods. oc, oral contraceptive.

| | Cohort | | Case-Control | | | |
|---|---|---|---|---|---|---|
| Pre-Menopause | Logistic Regression | | Logistic Regression | | Retrospective Likelihood | |
| | Estimate(SD) | P-value | Estimate(SD) | P-value | Estimate(SD) | P-value |
| oc 0-10 | -0.296(0.159) | 0.063 | -0.511(0.222) | 0.021 | -0.512(0.217) | 0.019 |
| oc >10 | 0.052(0.148) | 0.724 | -0.089(0.213) | 0.677 | -0.107(0.208) | 0.607 |
| oc 0-10 * PRS | 0.084(0.143) | 0.559 | 0.126(0.22) | 0.567 | 0.082(0.146) | 0.573 |
| oc >10 * PRS | -0.064(0.135) | 0.633 | 0.088(0.211) | 0.677 | -0.088(0.138) | 0.521 |
| | Estimate(SD) | P-value | Estimate(SD) | P-value | Estimate(SD) | P-value |
| oc 0-5 | -0.352(0.185) | 0.057 | -0.53(0.25) | 0.034 | -0.52(0.245) | 0.034 |
| oc 5-10 | -0.259(0.174) | 0.136 | -0.502(0.239) | 0.036 | -0.507(0.235) | 0.031 |
| oc 10-15 | -0.058(0.166) | 0.728 | -0.114(0.239) | 0.632 | -0.159(0.233) | 0.495 |
| oc >15 | 0.112(0.153) | 0.464 | -0.076(0.22) | 0.730 | -0.08(0.215) | 0.710 |
| oc 0-5 * PRS | 0.145(0.163) | 0.373 | 0.161(0.257) | 0.530 | 0.124(0.167) | 0.457 |
| oc 5-10 * PRS | 0.036(0.156) | 0.820 | 0.106(0.238) | 0.655 | 0.047(0.16) | 0.767 |
| oc 10-15 * PRS | -0.069(0.152) | 0.647 | 0.13(0.24) | 0.588 | -0.084(0.156) | 0.591 |
| oc >15 * PRS | -0.062(0.139) | 0.658 | 0.064(0.22) | 0.770 | -0.092(0.142) | 0.516 |

Table 4: Parameter estimates of the log odds ratio, standard deviation, and p-values for the disease-risk model in the UK Biobank breast cancer data for premenopausal women. Years of OC use is categorized into two sets: 1) never, 0-10 years, and $>$ 10 years. 2) never, 0-5 years, 5-10 years, 10-15 years, and $>$ 15 years. The reference group is women who have never taken OC. The model setting is the same as the continuous years of OC use model for all methods. oc, oral contraceptive.

| Post-Menopause | Estimate(SD) | P-value |
|---|---|---|
| eta_PC1 | -0.006(0.004) | 0.136 |
| eta_PC2 | -0.001(0.005) | 0.782 |
| eta_PC3 | -0.003(0.008) | 0.732 |
| eta_PC4 | -0.010(0.004) | 0.018 |
| eta_PC5 | -0.001(0.003) | 0.718 |
| eta_PC6 | 0.003(0.007) | 0.696 |
| eta_PC7 | 0.002(0.005) | 0.700 |
| eta_PC8 | 0.002(0.005) | 0.765 |
| eta_PC9 | 0.004(0.004) | 0.224 |
| eta_PC10 | 0.005(0.007) | 0.485 |
| eta_birth.x | -0.002(0.026) | 0.941 |
| eta_birth.y | -0.001(0.027) | 0.958 |
| sigma_11001 | 0.909(0.057) | - |
| sigma_10003 | 0.382(0.129) | - |
| sigma_11002 | 0.899(0.049) | - |
| sigma_11003 | 0.966(0.051) | - |
| sigma_11004 | 0.939(0.044) | - |
| sigma_11005 | 0.960(0.048) | - |
| sigma_11006 | 0.952(0.047) | - |
| sigma_11007 | 0.969(0.038) | - |
| sigma_11008 | 1.000(0.040) | - |
| sigma_11009 | 0.986(0.037) | - |
| sigma_11010 | 0.939(0.034) | - |
| sigma_11011 | 0.944(0.033) | - |
| sigma_11012 | 0.928(0.066) | - |
| sigma_11013 | 1.028(0.041) | - |
| sigma_11014 | 0.987(0.044) | - |
| sigma_11016 | 0.995(0.036) | - |
| sigma_11017 | 0.984(0.050) | - |
| sigma_11018 | 0.998(0.052) | - |
| sigma_11020 | 0.969(0.047) | - |
| sigma_11021 | 1.009(0.050) | - |
| sigma_11022 | 1.161(0.150) | - |
| sigma_11023 | 0.567(0.202) | - |

Table 5: The $\eta_s$ and $\sigma_{S^c}$ estimates from the PRS model of the retrospective likelihood method in the UK Biobank breast cancer data for postmenopausal women. Here we assumed independence between PRS and the risk factors conditional on $S =$ (Top 10 PCs, North coordinates of birth location, East coordinates of birth location) and stratification factor $S^c =$ (Assessment Centres).

| Pre-Menopause | Estimate(SD) | P-value |
|---|---|---|
| eta_PC1 | -0.006(0.007) | 0.389 |
| eta_PC2 | -0.0002(0.009) | 0.978 |
| eta_PC3 | 0.013(0.014) | 0.370 |
| eta_PC4 | 0.004(0.006) | 0.494 |
| eta_PC5 | -0.003(0.004) | 0.508 |
| eta_PC6 | 0.0002(0.015) | 0.990 |
| eta_PC7 | 0.008(0.007) | 0.209 |
| eta_PC8 | -0.002(0.010) | 0.851 |
| eta_PC9 | -0.0003(0.007) | 0.966 |
| eta_PC10 | -0.002(0.012) | 0.847 |
| eta_birth.x | 0.009(0.042) | 0.820 |
| eta_birth.y | 0.006(0.045) | 0.895 |
| sigma_11001 | 0.914(0.088) | - |
| sigma_10003 | 0.264(0.222) | - |
| sigma_11002 | 0.944(0.088) | - |
| sigma_11003 | 0.966(0.081) | - |
| sigma_11004 | 0.969(0.074) | - |
| sigma_11005 | 0.944(0.077) | - |
| sigma_11006 | 0.972(0.076) | - |
| sigma_11007 | 0.921(0.058) | - |
| sigma_11008 | 0.937(0.065) | - |
| sigma_11009 | 0.996(0.062) | - |
| sigma_11010 | 1.020(0.055) | - |
| sigma_11011 | 0.943(0.052) | - |
| sigma_11012 | 0.891(0.120) | - |
| sigma_11013 | 0.904(0.065) | - |
| sigma_11014 | 0.912(0.073) | - |
| sigma_11016 | 0.934(0.069) | - |
| sigma_11017 | 1.067(0.084) | - |
| sigma_11018 | 0.923(0.067) | - |
| sigma_11020 | 1.102(0.093) | - |
| sigma_11021 | 0.929(0.090) | - |
| sigma_11022 | 0.707(0.293) | - |

Table 6: The $\eta_s$ and $\sigma_{S^c}$ estimates from the PRS model of the retrospective likelihood method in the UK Biobank breast cancer data for premenopausal women. Here we assumed independence between PRS and the risk factors conditional on $S$ = (Top 10 PCs, North coordinates of birth location, East coordinates of birth location) and stratification factor $S^c$ = (Assessment Centres).

| Post-Menopause | Retrospective Likelihood | | Logistic Regression | | | |
|---|---|---|---|---|---|---|
| | | | Cohort | | Case-Control | |
| | Estimate(SD) | P-value | Estimate(SD) | P-value | Estimate(SD) | P-value |
| PRS | 0.557(0.196) | 0.005 | 0.548(0.194) | 0.005 | 0.463(0.249) | 0.063 |
| PC1 | -0.002(0.009) | 0.808 | -0.002(0.008) | 0.757 | -0.003(0.01) | 0.788 |
| PC2 | -0.009(0.011) | 0.418 | -0.012(0.007) | 0.100 | -0.01(0.011) | 0.382 |
| PC3 | -0.003(0.017) | 0.882 | -0.003(0.01) | 0.769 | -0.004(0.017) | 0.819 |
| PC4 | 0.002(0.009) | 0.787 | 0.002(0.006) | 0.697 | 0.003(0.009) | 0.766 |
| PC5 | -0.002(0.006) | 0.689 | 0.002(0.004) | 0.582 | -0.002(0.006) | 0.724 |
| PC6 | -0.041(0.016) | 0.010 | -0.018(0.012) | 0.145 | -0.04(0.016) | 0.012 |
| PC7 | -0.005(0.011) | 0.637 | -0.005(0.007) | 0.504 | -0.006(0.011) | 0.552 |
| PC8 | 0.02(0.012) | 0.101 | 0.01(0.009) | 0.249 | 0.019(0.012) | 0.128 |
| PC9 | -0.004(0.008) | 0.657 | -0.007(0.006) | 0.201 | -0.004(0.008) | 0.661 |
| PC10 | -0.018(0.016) | 0.255 | 0.001(0.011) | 0.942 | -0.021(0.016) | 0.196 |
| HRT.ever | 0.11(0.067) | 0.104 | 0.132(0.051) | 0.009 | 0.106(0.069) | 0.122 |
| agegroups[50,60) | 0.381(0.238) | 0.109 | 0.45(0.202) | 0.026 | 0.405(0.24) | 0.091 |
| agegroups[60,70) | 0.527(0.238) | 0.027 | 0.683(0.202) | 0.001 | 0.548(0.24) | 0.023 |
| agegroups[70,80) | 0.484(0.48) | 0.313 | 0.879(0.355) | 0.013 | 0.498(0.482) | 0.302 |
| n.children | -0.077(0.052) | 0.138 | -0.078(0.04) | 0.050 | -0.084(0.053) | 0.109 |
| birth.x | 0.021(0.046) | 0.649 | 0.031(0.033) | 0.350 | 0.026(0.047) | 0.578 |
| birth.y | 0.011(0.055) | 0.834 | 0.014(0.039) | 0.716 | 0.004(0.055) | 0.944 |
| age.firstbirth.group[25,35) | 0.016(0.077) | 0.840 | 0.014(0.059) | 0.817 | 0.016(0.079) | 0.835 |
| age.firstbirth.group[35,80) | 0.278(0.169) | 0.100 | 0.258(0.117) | 0.027 | 0.26(0.17) | 0.125 |
| age.firstbirth.group.never | -0.161(0.148) | 0.276 | -0.003(0.112) | 0.978 | -0.165(0.151) | 0.272 |
| height | 0.087(0.034) | 0.010 | 0.134(0.026) | 0.000 | 0.084(0.034) | 0.015 |
| age.menarche | 0.018(0.033) | 0.580 | -0.016(0.025) | 0.520 | 0.027(0.034) | 0.414 |
| age.menopause | 0.095(0.034) | 0.005 | 0.097(0.026) | 0.000 | 0.092(0.034) | 0.007 |
| years.oc | -0.008(0.034) | 0.819 | 0.026(0.025) | 0.306 | -0.003(0.034) | 0.934 |
| BMI | 0.138(0.034) | $4.37\times10^{-5}$ | 0.148(0.024) | $5.99\times10^{-10}$ | 0.141(0.034) | $4.28\times10^{-5}$ |
| assessment_center10003 | -0.986(0.955) | 0.302 | 0.042(0.732) | 0.954 | -0.908(0.973) | 0.351 |
| assessment_center11002 | -0.794(0.271) | 0.003 | -0.125(0.176) | 0.478 | -0.795(0.277) | 0.004 |
| assessment_center11003 | -0.57(0.284) | 0.045 | -0.359(0.181) | 0.047 | -0.573(0.291) | 0.049 |
| assessment_center11004 | -0.635(0.275) | 0.021 | -0.22(0.176) | 0.211 | -0.628(0.282) | 0.026 |
| assessment_center11005 | -0.55(0.277) | 0.047 | -0.208(0.173) | 0.229 | -0.566(0.284) | 0.046 |
| assessment_center11006 | -0.672(0.263) | 0.011 | -0.147(0.167) | 0.379 | -0.66(0.269) | 0.014 |
| assessment_center11007 | -0.499(0.249) | 0.045 | -0.129(0.153) | 0.397 | -0.51(0.254) | 0.044 |
| assessment_center11008 | -0.652(0.247) | 0.008 | -0.209(0.154) | 0.174 | -0.603(0.253) | 0.017 |
| assessment_center11009 | -0.584(0.249) | 0.019 | -0.301(0.153) | 0.050 | -0.557(0.254) | 0.028 |
| assessment_center11010 | -0.83(0.24) | 0.001 | -0.445(0.149) | 0.003 | -0.785(0.244) | 0.001 |
| assessment_center11011 | -0.687(0.242) | 0.005 | -0.339(0.15) | 0.024 | -0.65(0.247) | 0.009 |
| assessment_center11012 | -0.849(0.306) | 0.006 | -0.283(0.204) | 0.166 | -0.86(0.312) | 0.006 |
| assessment_center11013 | -0.776(0.247) | 0.002 | -0.406(0.154) | 0.008 | -0.806(0.252) | 0.001 |
| assessment_center11014 | -1.031(0.254) | $4.83\times10^{-5}$ | -0.684(0.164) | $3.05\times10^{-5}$ | -1.016(0.259) | $9.04\times10^{-5}$ |
| assessment_center11016 | -0.721(0.241) | 0.003 | -0.223(0.15) | 0.137 | -0.688(0.247) | 0.005 |
| assessment_center11017 | -0.876(0.27) | 0.001 | -0.474(0.173) | 0.006 | -0.873(0.275) | 0.001 |
| assessment_center11018 | -1.14(0.271) | $2.66\times10^{-5}$ | -0.727(0.179) | $5.11\times10^{-5}$ | -1.132(0.277) | $4.34\times10^{-5}$ |
| assessment_center11020 | -1.039(0.264) | $8.26\times10^{-5}$ | -0.546(0.172) | 0.001 | -1.026(0.269) | $1.34\times10^{-4}$ |
| assessment_center11021 | -0.923(0.263) | $4.56\times10^{-4}$ | -0.571(0.171) | 0.001 | -0.977(0.269) | $2.82\times10^{-4}$ |
| assessment_center11022 | -0.768(0.501) | 0.125 | -0.296(0.338) | 0.381 | -0.863(0.525) | 0.100 |
| assessment_center11023 | -13.88(430.057) | 0.974 | -12.204(128.064) | 0.924 | -13.88(251.871) | 0.956 |
| alcohol | 0.235(0.08) | 0.003 | 0.194(0.064) | 0.002 | 0.236(0.082) | 0.004 |
| PRS*HRT.ever | -0.023(0.047) | 0.624 | -0.01(0.047) | 0.838 | -0.048(0.072) | 0.506 |
| PRS*birth.x | -0.079(0.037) | 0.035 | -0.068(0.025) | 0.006 | -0.09(0.04) | 0.024 |
| PRS*birth.y | -0.019(0.038) | 0.612 | -0.012(0.025) | 0.632 | -0.014(0.041) | 0.727 |
| PRS*years.oc | -0.05(0.024) | 0.037 | -0.055(0.023) | 0.019 | -0.032(0.036) | 0.373 |
| PRS*age.menopause | $2.00\times10^{-5}$(0.024) | 0.999 | -0.002(0.024) | 0.933 | -0.015(0.036) | 0.678 |
| PRS*age.menarche | -0.014(0.023) | 0.561 | -0.01(0.023) | 0.673 | 0.018(0.036) | 0.613 |
| PRS*BMI | -0.012(0.023) | 0.609 | -0.01(0.022) | 0.663 | 0.023(0.037) | 0.524 |
| PRS*agegroups[50,60) | -0.022(0.188) | 0.907 | -0.05(0.188) | 0.788 | 0.136(0.243) | 0.575 |
| PRS*agegroups[60,70) | -0.037(0.188) | 0.842 | -0.055(0.188) | 0.771 | 0.12(0.243) | 0.622 |
| PRS*agegroups[70,80) | -0.547(0.347) | 0.115 | -0.545(0.35) | 0.120 | -0.791(0.512) | 0.122 |
| PRS*alcohol | -0.099(0.058) | 0.086 | -0.091(0.057) | 0.112 | -0.121(0.086) | 0.160 |
| PRS*height | -0.013(0.024) | 0.593 | -0.02(0.023) | 0.383 | -0.031(0.036) | 0.396 |
| PRS*n.children | 0.011(0.037) | 0.758 | 0.005(0.036) | 0.883 | -0.036(0.056) | 0.522 |
| PRS*age.firstbirth.group[25,35) | 0.084(0.054) | 0.123 | 0.108(0.054) | 0.045 | 0.083(0.082) | 0.309 |
| PRS*age.firstbirth.group[35,80) | -0.14(0.114) | 0.219 | -0.083(0.111) | 0.454 | -0.372(0.17) | 0.029 |
| PRS*age.firstbirth.group.never | -0.034(0.105) | 0.750 | -0.019(0.103) | 0.854 | -0.122(0.159) | 0.445 |

Table 7: Parameter estimates of the log odds ratio, standard deviation, and p-values for the disease-risk model in the UK Biobank breast cancer data for postmenopausal women. PRS, breast cancer-related polygenic risk score constructed from 313 SNPs; Birth.x, the East coordinates of individual's birth location; Birth.y, the North coordinates of individual's birth location; oc, oral contraceptive.

| Pre-Menopause | Retrospective Likelihood | | Logistic Regression | | | |
|---|---|---|---|---|---|---|
| | | | Cohort | | Case-Control | |
| | Estimate(SD) | P-value | Estimate(SD) | P-value | Estimate(SD) | P-value |
| PRS | 0.454(0.164) | 0.006 | 0.434(0.158) | 0.006 | 0.602(0.243) | 0.013 |
| PC1 | -0.026(0.023) | 0.274 | -0.024(0.017) | 0.151 | -0.029(0.025) | 0.241 |
| PC2 | -0.035(0.025) | 0.157 | -0.02(0.014) | 0.168 | -0.038(0.025) | 0.137 |
| PC3 | 0.039(0.031) | 0.215 | 0.012(0.017) | 0.467 | 0.041(0.032) | 0.198 |
| PC4 | 0.004(0.015) | 0.809 | -0.009(0.009) | 0.314 | 0.004(0.015) | 0.816 |
| PC5 | -0.007(0.009) | 0.461 | -0.001(0.006) | 0.889 | -0.008(0.009) | 0.409 |
| PC6 | 0.07(0.033) | 0.034 | 0.048(0.021) | 0.023 | 0.065(0.034) | 0.054 |
| PC7 | 0.024(0.015) | 0.111 | 0.019(0.01) | 0.065 | 0.023(0.015) | 0.122 |
| PC8 | -0.06(0.022) | 0.005 | -0.025(0.014) | 0.081 | -0.056(0.022) | 0.011 |
| PC9 | -0.001(0.015) | 0.969 | 0.001(0.01) | 0.926 | -0.001(0.015) | 0.971 |
| PC10 | -0.019(0.027) | 0.483 | 0.009(0.017) | 0.619 | -0.022(0.028) | 0.430 |
| agegroups[45,50) | 0.365(0.128) | 0.004 | 0.193(0.096) | 0.043 | 0.333(0.131) | 0.011 |
| agegroups[50,80) | 0.251(0.156) | 0.108 | 0.156(0.119) | 0.190 | 0.278(0.162) | 0.086 |
| n.children | 0.166(0.099) | 0.095 | 0.131(0.076) | 0.085 | 0.13(0.101) | 0.197 |
| birth.x | 0.046(0.079) | 0.557 | 0.006(0.057) | 0.911 | 0.055(0.081) | 0.499 |
| birth.y | 0.071(0.092) | 0.438 | 0.081(0.065) | 0.216 | 0.071(0.094) | 0.448 |
| age.firstbirth.group[25,35) | 0.214(0.166) | 0.198 | 0.302(0.129) | 0.019 | 0.24(0.169) | 0.156 |
| age.firstbirth.group[35,80) | 0.542(0.244) | 0.026 | 0.518(0.184) | 0.005 | 0.562(0.251) | 0.025 |
| age.firstbirth.group.never | 0.756(0.275) | 0.006 | 0.695(0.212) | 0.001 | 0.687(0.281) | 0.014 |
| height | 0.08(0.058) | 0.168 | 0.105(0.044) | 0.016 | 0.078(0.059) | 0.186 |
| age.menarche | -0.029(0.057) | 0.617 | -0.007(0.043) | 0.868 | -0.015(0.058) | 0.793 |
| years.oc | 0.085(0.057) | 0.135 | 0.143(0.041) | 0.001 | 0.095(0.059) | 0.106 |
| BMI | 0.039(0.058) | 0.501 | 0.019(0.044) | 0.665 | 0.042(0.059) | 0.478 |
| assessment_center10003 | 14.416(1352.772) | 0.991 | 0.441(1.041) | 0.672 | 14.416(882.744) | 0.987 |
| assessment_center11002 | 0.276(0.444) | 0.534 | 0.168(0.302) | 0.577 | 0.301(0.461) | 0.514 |
| assessment_center11003 | 0.666(0.443) | 0.132 | 0.337(0.294) | 0.252 | 0.613(0.457) | 0.180 |
| assessment_center11004 | 0.193(0.433) | 0.656 | 0.127(0.299) | 0.670 | 0.222(0.451) | 0.622 |
| assessment_center11005 | 0.228(0.427) | 0.592 | 0.068(0.293) | 0.816 | 0.217(0.442) | 0.624 |
| assessment_center11006 | 0.362(0.406) | 0.373 | 0.41(0.281) | 0.144 | 0.264(0.421) | 0.531 |
| assessment_center11007 | 0.074(0.377) | 0.844 | 0.245(0.267) | 0.357 | 0.145(0.392) | 0.712 |
| assessment_center11008 | 0.437(0.387) | 0.259 | 0.388(0.266) | 0.145 | 0.62(0.404) | 0.125 |
| assessment_center11009 | 0.282(0.383) | 0.462 | 0.133(0.266) | 0.618 | 0.231(0.399) | 0.564 |
| assessment_center11010 | 0.287(0.36) | 0.424 | 0.119(0.253) | 0.638 | 0.286(0.375) | 0.445 |
| assessment_center11011 | -0.11(0.368) | 0.765 | -0.084(0.263) | 0.749 | -0.065(0.384) | 0.866 |
| assessment_center11012 | -0.21(0.539) | 0.698 | -0.541(0.393) | 0.168 | -0.203(0.558) | 0.716 |
| assessment_center11013 | 0.292(0.394) | 0.459 | -0.077(0.271) | 0.776 | 0.149(0.409) | 0.716 |
| assessment_center11014 | -0.046(0.403) | 0.909 | -0.283(0.288) | 0.327 | -0.036(0.418) | 0.932 |
| assessment_center11016 | -0.147(0.395) | 0.709 | -0.245(0.286) | 0.392 | -0.075(0.409) | 0.855 |
| assessment_center11017 | -0.593(0.415) | 0.153 | -0.191(0.309) | 0.537 | -0.655(0.433) | 0.131 |
| assessment_center11018 | -0.301(0.394) | 0.445 | -0.001(0.286) | 0.998 | -0.396(0.409) | 0.334 |
| assessment_center11020 | -0.456(0.428) | 0.286 | -0.342(0.312) | 0.273 | -0.49(0.444) | 0.270 |
| assessment_center11021 | -0.167(0.443) | 0.706 | -0.406(0.318) | 0.201 | -0.265(0.457) | 0.562 |
| assessment_center11022 | -14.223(908.653) | 0.988 | -12.367(193.462) | 0.949 | -14.223(490.004) | 0.977 |
| assessment_center11023 | - | - | -12.288(339.894) | 0.971 | - | - |
| alcohol | 0.068(0.163) | 0.675 | 0.176(0.124) | 0.155 | 0.067(0.167) | 0.689 |
| PRS*birth.x | -0.021(0.063) | 0.740 | -0.015(0.044) | 0.740 | -0.027(0.071) | 0.699 |
| PRS*birth.y | 0.003(0.063) | 0.967 | -0.003(0.042) | 0.946 | -0.006(0.07) | 0.932 |
| PRS*years.oc | -0.084(0.039) | 0.034 | -0.081(0.037) | 0.031 | -0.014(0.062) | 0.824 |
| PRS*age.menarche | -0.025(0.04) | 0.526 | -0.032(0.04) | 0.428 | 0.016(0.064) | 0.800 |
| PRS*BMI | -0.032(0.04) | 0.417 | -0.03(0.041) | 0.464 | -0.012(0.064) | 0.852 |
| PRS*agegroups[45,50) | 0.013(0.091) | 0.885 | 0.029(0.087) | 0.737 | -0.096(0.141) | 0.494 |
| PRS*agegroups[50,80) | 0.126(0.11) | 0.255 | 0.146(0.106) | 0.169 | 0.212(0.174) | 0.223 |
| PRS*alcohol | 0.069(0.118) | 0.554 | 0.057(0.114) | 0.615 | -0.058(0.187) | 0.758 |
| PRS*height | -0.004(0.04) | 0.928 | 0.001(0.039) | 0.981 | 0.05(0.063) | 0.422 |
| PRS*n.children | 0.015(0.07) | 0.832 | -0.004(0.068) | 0.959 | -0.102(0.11) | 0.355 |
| PRS*age.firstbirth.group[25,35) | -0.014(0.119) | 0.908 | -0.041(0.115) | 0.722 | 0.103(0.177) | 0.561 |
| PRS*age.firstbirth.group[35,80) | 0.04(0.168) | 0.813 | 0.025(0.166) | 0.881 | 0.118(0.255) | 0.643 |
| PRS*age.firstbirth.group.never | -0.0005(0.195) | 0.998 | -0.057(0.191) | 0.765 | -0.215(0.295) | 0.465 |

Table 8: Parameter estimates of the log odds ratio, standard deviation, and p-values for the disease-risk model in the UK Biobank breast cancer data for premenopausal women. PRS, breast cancer-related polygenic risk score constructed from 313 SNPs; Birth.x, the East co-ordinates of individual's birth location; Birth.y, the North co-ordinates of individual's birth location; oc, oral contraceptive.

| Colorectal Cancer | Estimate(SD) | P-value |
|---|---|---|
| eta_PC1 | 0.007(0.007) | 0.363 |
| eta_PC2 | -0.006(0.009) | 0.492 |
| eta_PC3 | 0.002(0.010) | 0.865 |
| eta_PC4 | 0.0002(0.005) | 0.969 |
| eta_PC5 | -0.002(0.003) | 0.467 |
| eta_PC6 | -0.0002(0.010) | 0.981 |
| eta_PC7 | 0.010(0.005) | 0.083 |
| eta_PC8 | -0.0005(0.007) | 0.944 |
| eta_PC9 | -0.001(0.004) | 0.758 |
| eta_PC10 | 0.005(0.008) | 0.528 |
| eta_birth.x | -0.006(0.025) | 0.822 |
| eta_birth.y | 0.020(0.026) | 0.453 |
| sigma_11001 | 0.982(0.061) | - |
| sigma_11002 | 0.944(0.056) | - |
| sigma_11003 | 1.006(0.055) | - |
| sigma_11004 | 0.979(0.053) | - |
| sigma_11005 | 0.954(0.053) | - |
| sigma_11006 | 0.907(0.051) | - |
| sigma_11007 | 0.984(0.042) | - |
| sigma_11008 | 0.958(0.042) | - |
| sigma_11009 | 0.964(0.038) | - |
| sigma_11010 | 0.987(0.036) | - |
| sigma_11011 | 0.964(0.036) | - |
| sigma_11012 | 0.997(0.075) | - |
| sigma_11013 | 1.030(0.043) | - |
| sigma_11014 | 0.893(0.040) | - |
| sigma_11016 | 0.979(0.040) | - |
| sigma_11017 | 0.873(0.051) | - |
| sigma_11018 | 0.964(0.051) | - |
| sigma_11020 | 1.024(0.056) | - |
| sigma_11021 | 1.025(0.056) | - |
| sigma_11022 | 0.847(0.170) | - |
| sigma_11023 | 1.334(0.565) | - |

Table 9: The $\eta_s$ and $\sigma_{S^c}$ estimates from the PRS model of the retrospective likelihood method in the UK Biobank colorectal cancer data. Here we assumed independence between PRS and the risk factors conditional on $S$ = (Top 10 PCs, North coordinates of birth location, East coordinates of birth location) and stratification factor $S^c$ = (Assessment Centres).

| | | | Logistic Regression | | | |
|---|---|---|---|---|---|---|
| Colorectal Cancer | Retrospective Likelihood | | Cohort | | Case-Control | |
| | Estimate(SD) | P-value | Estimate(SD) | P-value | Estimate(SD) | P-value |
| PRS | 0.749(0.196) | $1.32\times10^{-4}$ | 0.655(0.186) | $4.12\times10^{-4}$ | 0.499(0.239) | 0.036 |
| PC1 | -0.033(0.018) | 0.064 | -0.024(0.012) | 0.041 | -0.031(0.017) | 0.072 |
| PC2 | -0.003(0.021) | 0.900 | 0.014(0.014) | 0.321 | -0.002(0.021) | 0.937 |
| PC3 | 0.002(0.022) | 0.917 | 0.007(0.015) | 0.618 | 0.001(0.022) | 0.948 |
| PC4 | -0.008(0.011) | 0.458 | -0.004(0.007) | 0.617 | -0.008(0.011) | 0.486 |
| PC5 | 0.012(0.006) | 0.055 | 0.01(0.004) | 0.017 | 0.013(0.006) | 0.049 |
| PC6 | 0.004(0.021) | 0.835 | 0.003(0.014) | 0.834 | 0.004(0.022) | 0.855 |
| PC7 | -0.005(0.013) | 0.671 | 0.004(0.009) | 0.686 | -0.007(0.013) | 0.594 |
| PC8 | -0.036(0.016) | 0.021 | -0.027(0.011) | 0.012 | -0.037(0.016) | 0.019 |
| PC9 | -0.018(0.009) | 0.046 | -0.007(0.006) | 0.243 | -0.017(0.009) | 0.058 |
| PC10 | -0.013(0.018) | 0.469 | -0.013(0.012) | 0.263 | -0.01(0.018) | 0.569 |
| Activity-Moderate | -0.076(0.104) | 0.461 | 0.024(0.074) | 0.748 | -0.07(0.105) | 0.506 |
| Activity-High | -0.067(0.106) | 0.528 | 0.018(0.076) | 0.814 | -0.044(0.107) | 0.679 |
| age[45,50) | 0.647(0.237) | 0.006 | 0.71(0.228) | 0.002 | 0.677(0.239) | 0.005 |
| age[50,55) | 1.274(0.221) | $7.64\times10^{-9}$ | 1.383(0.21) | $4.46\times10^{-11}$ | 1.263(0.222) | $1.25\times10^{-8}$ |
| age[55,60) | 1.714(0.216) | $2.37\times10^{-15}$ | 1.651(0.205) | $8.51\times10^{-16}$ | 1.721(0.218) | $2.86\times10^{-15}$ |
| age[60,65) | 1.927(0.212) | $8.28\times10^{-20}$ | 1.896(0.201) | $4.56\times10^{-21}$ | 1.947(0.213) | $6.49\times10^{-20}$ |
| age[65,70) | 2.212(0.215) | $6.32\times10^{-25}$ | 2.149(0.202) | $1.98\times10^{-26}$ | 2.229(0.216) | $6.00\times10^{-25}$ |
| age[70,75) | 2.232(0.509) | $1.15\times10^{-5}$ | 2.398(0.330) | $3.86\times10^{-13}$ | 2.228(0.527) | $2.36\times10^{-5}$ |
| sexMale | 0.200(0.116) | 0.083 | 0.227(0.082) | 0.006 | 0.218(0.118) | 0.065 |
| alcohol | 0.017(0.103) | 0.871 | -0.015(0.075) | 0.837 | 0.023(0.105) | 0.828 |
| height | 0.038(0.054) | 0.479 | 0.057(0.039) | 0.140 | 0.035(0.056) | 0.534 |
| assessment_center10003 | - | - | -6.432(98.389) | 0.948 | - | - |
| assessment_center11002 | -0.169(0.292) | 0.563 | -0.039(0.184) | 0.834 | -0.189(0.298) | 0.526 |
| assessment_center11003 | -0.358(0.288) | 0.213 | -0.261(0.185) | 0.159 | -0.329(0.294) | 0.262 |
| assessment_center11004 | -0.122(0.293) | 0.677 | -0.296(0.185) | 0.111 | -0.155(0.302) | 0.607 |
| assessment_center11005 | -0.0004(0.291) | 0.999 | -0.171(0.183) | 0.349 | 0.004(0.299) | 0.989 |
| assessment_center11006 | -0.242(0.281) | 0.389 | -0.259(0.177) | 0.144 | -0.235(0.287) | 0.414 |
| assessment_center11007 | -0.175(0.26) | 0.502 | -0.169(0.162) | 0.299 | -0.163(0.266) | 0.540 |
| assessment_center11008 | -0.318(0.256) | 0.215 | -0.233(0.162) | 0.150 | -0.361(0.261) | 0.167 |
| assessment_center11009 | -0.317(0.252) | 0.209 | -0.273(0.161) | 0.089 | -0.328(0.259) | 0.206 |
| assessment_center11010 | -0.384(0.244) | 0.117 | -0.292(0.154) | 0.058 | -0.411(0.250) | 0.099 |
| assessment_center11011 | -0.215(0.251) | 0.391 | -0.241(0.156) | 0.122 | -0.228(0.256) | 0.373 |
| assessment_center11012 | -0.318(0.333) | 0.339 | -0.208(0.221) | 0.346 | -0.312(0.340) | 0.359 |
| assessment_center11013 | -0.361(0.257) | 0.160 | -0.347(0.162) | 0.032 | -0.412(0.262) | 0.117 |
| assessment_center11014 | -0.297(0.261) | 0.254 | -0.35(0.165) | 0.034 | -0.335(0.266) | 0.208 |
| assessment_center11016 | -0.510(0.251) | 0.042 | -0.318(0.159) | 0.045 | -0.583(0.256) | 0.023 |
| assessment_center11017 | -0.257(0.287) | 0.371 | -0.443(0.183) | 0.016 | -0.258(0.293) | 0.379 |
| assessment_center11018 | -0.334(0.279) | 0.231 | -0.36(0.178) | 0.044 | -0.377(0.285) | 0.187 |
| assessment_center11020 | -0.518(0.282) | 0.066 | -0.449(0.183) | 0.014 | -0.600(0.288) | 0.037 |
| assessment_center11021 | -0.738(0.281) | 0.009 | -0.687(0.187) | $2.42\times10^{-4}$ | -0.713(0.288) | 0.013 |
| assessment_center11022 | -1.070(0.659) | 0.105 | -0.953(0.472) | 0.043 | -0.932(0.675) | 0.167 |
| assessment_center11023 | -1.438(1.463) | 0.326 | -1.339(1.012) | 0.186 | -1.890(1.476) | 0.200 |
| waist | 0.117(0.044) | 0.008 | 0.130(0.031) | $2.16\times10^{-5}$ | 0.123(0.045) | 0.006 |
| smoke.status.ever | 0.168(0.079) | 0.033 | 0.173(0.056) | 0.002 | 0.180(0.080) | 0.025 |
| smoke.status.current | -0.087(0.134) | 0.515 | -0.084(0.101) | 0.402 | -0.099(0.136) | 0.467 |
| redmeat.numeric | 0.057(0.039) | 0.147 | 0.073(0.028) | 0.009 | 0.065(0.040) | 0.103 |
| process_meat | 0.102(0.041) | 0.013 | 0.067(0.029) | 0.023 | 0.099(0.042) | 0.017 |
| veg.numeric | -0.012(0.037) | 0.756 | 0.006(0.026) | 0.823 | -0.015(0.038) | 0.699 |
| birth.x | -0.039(0.042) | 0.363 | -0.057(0.029) | 0.047 | -0.036(0.043) | 0.397 |
| birth.y | -0.005(0.052) | 0.927 | 0.004(0.035) | 0.901 | -0.005(0.053) | 0.919 |
| PRS*birth.x | -0.004(0.036) | 0.913 | 0.006(0.023) | 0.799 | 0.019(0.039) | 0.616 |
| PRS*birth.y | 0.031(0.037) | 0.409 | 0.025(0.023) | 0.290 | 0.036(0.039) | 0.355 |
| PRS*Activity-Moderate | -0.132(0.069) | 0.054 | -0.117(0.066) | 0.076 | -0.091(0.109) | 0.401 |
| PRS*Activity-High | -0.084(0.07) | 0.230 | -0.074(0.067) | 0.271 | -0.011(0.112) | 0.919 |
| PRS*age[45,50) | -0.127(0.205) | 0.537 | -0.096(0.198) | 0.628 | -0.017(0.242) | 0.945 |
| PRS*age[50,55) | -0.123(0.188) | 0.512 | -0.109(0.182) | 0.550 | -0.175(0.223) | 0.431 |
| PRS*age[55,60) | -0.052(0.183) | 0.777 | -0.036(0.177) | 0.839 | 0.063(0.221) | 0.776 |
| PRS*age[60,65) | -0.172(0.180) | 0.338 | -0.134(0.174) | 0.440 | -0.13(0.212) | 0.542 |
| PRS*age[65,70) | -0.192(0.181) | 0.288 | -0.159(0.174) | 0.361 | -0.113(0.216) | 0.600 |
| PRS*age[70,75) | -0.969(0.314) | 0.002 | -0.801(0.294) | 0.007 | -0.875(0.507) | 0.084 |
| PRS*sexMale | 0.009(0.078) | 0.912 | 0.018(0.074) | 0.804 | 0.129(0.125) | 0.303 |
| PRS*alcohol | -0.071(0.070) | 0.313 | -0.057(0.067) | 0.394 | -0.002(0.107) | 0.986 |
| PRS*height | 0.018(0.036) | 0.624 | 0.017(0.035) | 0.622 | -0.041(0.059) | 0.487 |
| PRS*waist | -0.012(0.029) | 0.685 | -0.020(0.028) | 0.470 | -0.016(0.046) | 0.736 |
| PRS*veg.numeric | -0.014(0.026) | 0.575 | -0.011(0.024) | 0.641 | -0.019(0.037) | 0.604 |
| PRS*redmeat.numeric | -0.014(0.027) | 0.596 | -0.009(0.026) | 0.734 | 0.024(0.042) | 0.564 |
| PRS*smoke.status.ever | 0.019(0.053) | 0.720 | 0.021(0.051) | 0.675 | 0.124(0.086) | 0.147 |
| PRS*smoke.status.current | 0.065(0.093) | 0.485 | 0.063(0.090) | 0.481 | 0.121(0.137) | 0.378 |
| PRS*process_meat | -0.033(0.027) | 0.232 | -0.030(0.026) | 0.255 | -0.044(0.043) | 0.310 |

Table 10: Parameter estimates of the log odds ratio, standard deviation, and p-values for the disease-risk model in the UK Biobank colorectal cancer data. PRS, colorectal cancer related polygenic risk score constructed from 95 SNPs; Birth.x, the East co-ordinates of individual's birth location; Birth.y, the North co-ordinates of individual's birth location.

Kapoor, P. M., N. Mavaddat, P. P. Choudhury, A. N. Wilcox, S. Lindström, S. Behrens, K. Michailidou, J. Dennis, M. K. Bolla, Q. Wang, A. Jung, Z. Abu-Ful, T. Ahearn, I. L. Andrulis, H. Anton-Culver, V. Arndt, K. J. Aronson, P. L. Auer, L. E. B. Freeman, H. Becher, M. W. Beckmann, A. Beeghly-Fadiel, J. Benitez, L. Bernstein, S. E. Bojesen, H. Brauch, H. Brenner, T. Brüning, Q. Cai, D. Campa, F. Canzian, A. Carracedo, B. D. Carter, J. E. Castelao, S. J. Chanock, N. Chatterjee, G. Chenevix-Trench, C. L. Clarke, F. J. Couch, A. Cox, S. S. Cross, K. Czene, J. Y. Dai, H. S. Earp, A. B. Ekici, A. H. Eliassen, M. Eriksson, D. G. Evans, P. A. Fasching, J. Figueroa, L. Fritschi, M. Gabrielson, M. Gago-Dominguez, C. Gao, S. M. Gapstur, M. M. Gaudet, G. G. Giles, A. González-Neira, P. Guénel, L. Haeberle, C. A. Haiman, N. Håkansson, P. Hall, U. Hamann, S. Hatse, J. Heyworth, B. Holleczek, R. N. Hoover, J. L. Hopper, A. Howell, D. J. Hunter, A. Investigators, kConFab/AOCS Investigators, E. M. John, M. E. Jones, R. Kaaks, R. Keeman, C. M. Kitahara, Y.-D. Ko, S. Koutros, A. W. Kurian, D. Lambrechts, L. Le Marchand, E. Lee, F. Lejbkowicz, M. Linet, J. Lissowska, A. Llaneza, R. J. MacInnis, M. E. Martinez, T. Maurer, C. McLean, S. L. Neuhausen, W. G. Newman, A. Norman, K. M. O'Brien, A. F. Olshan, J. E. Olson, H. Olsson, N. Orr, C. M. Perou, G. Pita, E. C. Polley, R. L. Prentice, G. Rennert, H. S. Rennert, K. J. Ruddy, D. P. Sandler, C. Saunders, M. J. Schoemaker, B. Schöttker, F. Schumacher, C. Scott, R. J. Scott, X.-O. Shu, A. Smeets, M. C. Southey, J. J. Spinelli, J. Stone, A. J. Swerdlow, R. M. Tamimi, J. A. Taylor, M. A. Troester, C. M. Vachon, E. M. van Veen, X. Wang, C. R. Weinberg, C. Weltens, W. Willett, S. J. Winham, A. Wolk, X. R. Yang, W. Zheng, A. Ziogas, A. M. Dunning, P. D. P. Pharoah, M. K. Schmidt, P. Kraft, D. F. Easton, R. L. Milne, M. García-Closas, and J. Chang-Claude (2020). Combined Associations of a Polygenic Risk Score and Classical Risk Factors With Breast Cancer Risk. *JNCI: Journal of the National Cancer Institute 113*(3), 329–337.

Lambert, S. A., L. Gil, S. Jupp, S. C. Ritchie, Y. Xu, A. Buniello, A. McMahon, G. Abraham, M. Chapman, H. Parkinson, et al. (2021). The polygenic score catalog as an open database for reproducibility and systematic evaluation. *Nature Genetics 53*(4), 420–425.

Li, H., M. H. Gail, S. Berndt, and N. Chatterjee (2010). Using cases to strengthen inference on the association between single nucleotide polymorphisms and a secondary phenotype in genome-wide association studies. *Genetic epidemiology 34*(5), 427–433.

Mavaddat, N., K. Michailidou, J. Dennis, M. Lush, L. Fachal, A. Lee, J. P. Tyrer, T.-H. Chen, Q. Wang, M. K. Bolla, et al. (2019). Polygenic risk scores for prediction of breast cancer and breast cancer subtypes. *The American Journal of Human Genetics 104*(1), 21–34.

Pal Choudhury, P., A. N. Wilcox, M. N. Brook, Y. Zhang, T. Ahearn, N. Orr, P. Coulson, M. J. Schoemaker, M. E. Jones, M. H. Gail, A. J. Swerdlow, N. Chatterjee, and M. Garcia-Closas (2019). Comparative Validation of Breast Cancer Risk Prediction Models and Projections for Future Risk Stratification. *JNCI: Journal of the National Cancer Institute 112*(3), 278–285.

PGS Catalog Team (2022). The polygenic score catalog calculator. Available at `https://github.com/PGScatalog/pgsc_calc`, version 1.3.0.
